# Localization and function of septins are susceptible to epitope tagging

**DOI:** 10.1101/2025.05.13.653749

**Authors:** Jack R. Gregory, Ian Mikale A. Llaneza, Aysha H. Osmani, Haley E. Gosselin, S. Amirreza Sabzian, Jian-Qiu Wu

## Abstract

Septins are hetero-oligomeric cytoskeletal proteins that assemble into filaments and scaffolds on the plasma membrane to aid cytokinesis, morphogenesis, and other cellular processes. Epitope tagging is widely used to study septin localization and function. However, functionality testing of tagged septins is often insufficient because of technical challenges. Fission yeast provides an ideal genetic system to test functionalities and localizations of tagged septins. mEGFP/mYFP tagged septins Spn1 and Spn4 localize exclusively to the division site as double rings during cytokinesis, but tdTomato tagged septins also localize to puncta or short linear structures across the plasma membrane. It was proposed that these additional septin structures serve as diffusion barriers and are important for the localizations and functions of several proteins, including the NDR-kinase Sid2 and active Cdc42 GTPase. By analyzing cell morphology, cytokinesis defects, and genetic interactions between tagged septins and three mutations, we find that septins are less functional with tdTomato or 3HA than other tags. Additionally, Sid2 appearance at the division site is after septins and delayed in septin deletions, contrary to previous reports. Our data re-emphasize the need for rigorous functional tests of tagged septins and for caution in interpreting function and localization data when using epitope tagged septins.

**SIGNIFICANCE STATEMENT:**

- Fission yeast septins Spn1 and Spn4, expressed under their native promoters, have drastically different localizations depending on the fluorescent tag used.
- By assessing cell morphology, cytokinesis defects, cell integrity, and genetic interactions between tagged septins and other mutations, we find that septins are less functional when tagged with tdTomato/3HA compared to mEGFP/mYFP.
- Our data highlight the need for rigorous functional tests of tagged septins and caution when interpreting localization/function data because septin polymers are susceptible to perturbations by epitope tags. Our results also caution the caveats of using epitope tags to study other proteins, which is almost indispensable.

## INTRODUCTION

Septins are a group of proteins considered to be the fourth element of the cytoskeleton (Mostowy and Cossart, 2012). The mutants affecting septins were first isolated in a landmark screen for cell division regulatory proteins in the budding yeast *Saccharomyces cerevisiae* (Hartwell, 1971). Subsequent pioneering experiments, including cloning and immunofluorescence staining, by the Pringle laboratory revealed that Cdc3, Cdc10, Cdc11, and Cdc12 localize to the cell-division site and are essential for cytokinesis and morphogenesis, among other processes (Haarer and Pringle, 1987; Ford and Pringle, 1991; Kim *et al*., 1991). This family of proteins were subsequently given their name, Septins, and found to exist in most eukaryotic cells; the exceptions are amoebozoa, excavate, land plants, and a few members of other eukaryotic supergroups where septins were independently lost (Neufeld and Rubin, 1994; Kinoshita *et al*., 1997; Nguyen *et al*., 2000; Surka *et al*., 2002; Pan *et al*., 2007; Kwitny *et al*., 2010; Juvvadi *et al*., 2011; Nishihama *et al*., 2011; Yamazaki *et al*., 2013; Onishi and Pringle, 2016; Delic *et al*., 2024).

Septins are essential for cytokinesis, cell polarization, exocytosis, neuronal development, and many other cellular processes in eukaryotic cells. They act as a membrane barrier to diffusion between the mother cell and the emerging bud in budding yeast (Takizawa *et al*., 2000) and as membrane diffusion barriers in mouse sperm and cilia (Kwitny *et al*., 2010). Budding yeast septins function as a scaffold on the plasma membrane to coordinate contractile-ring assembly, the synthesis and deposition of chitin in new cell walls, and most other processes at the bud neck (Gladfelter *et al*., 2001). Septins also serve as membrane curvature sensors, helping configure and compartmentalize the plasma membrane (Bridges *et al*., 2016; Curtis *et al*., 2025; Edelmaier *et al*., 2025). In filamentous fungi, including fungal pathogens, septins form various distinct structures within cells which are important for fungal morphogenesis and infection (Gladfelter, 2006; Harris, 2006). Septins bind and stabilize microtubules and are required for the formation of neurites (Surka *et al*., 2002). In fission yeast and neurons, septins have been implicated in vesicle trafficking pathways and shown to interact with the exocyst complex during cytokinesis (Singh *et al*., 2024). Considering their numerous functions, it is unsurprising that mutations in septins have been implicated in many diseases, including Alzheimer’s Disease, cancer, bipolar disorder, infertility, birth defects, and other neurological disorders (Kinoshita *et al*., 1998; Pissuti Damalio *et al*., 2012; Das and Kunwar, 2025). It has also been shown that septins are essential for host defense against a variety of viral infections, including hepatitis C, influenza, and vaccinia (Khairat *et al*., 2024). Moreover, due to the essential roles of septins in membrane transport in various fungal species, septins are potential targets for anti-fungal drug development (Eisermann *et al*., 2023). Thus, it is crucial to have a precise understanding of how septins localize and function throughout the cell cycle to allow for accurate investigation of the role of septins in human disease.

Septins are GTP binding proteins, and most cell types express several different septins. Septins are capable of binding to one another to form multimeric complexes, often as hexamers or octamers (Garcia *et al*., 2011; Fung *et al*., 2014; Cavini *et al*., 2021). These complexes can further polymerize into various higher-order structures, such as rings, hourglasses, filaments, bars, patches, and gauzes (Byers and Goetsch, 1976; Rodal *et al*., 2005; Vrabioiu and Mitchison, 2006; Garcia *et al*., 2011; Bridges *et al*., 2014; El Alaoui *et al*., 2025). Depending on their function, septin structures occupy different locations in cells, including cilia, division sites, associations with microtubules, and penetrating structures of fungal pathogens (Bridges and Gladfelter, 2015). The most notable and well-studied location of septins is at the cell-division site, where they are crucial or essential for cytokinesis (Haarer and Pringle, 1987; Ford and Pringle, 1991; Kim *et al*., 1991). A structure of septins, determined via electron microscopy (Sirajuddin *et al*., 2007), shows that septins interact via two faces: the G interface (GTP binding domain) and the NC interface (N- and C-termini). Most septins also contain a septin unique element (SUE) and a coiled-coil domain, most commonly within their C-termini (Sirajuddin *et al*., 2007; Shuman and Momany, 2021). Upstream of the GTP binding domain is a polybasic region, which contains many basic amino acids that interact with phosphatidylinositol phosphates on the plasma membrane (Bridges and Gladfelter, 2015). Septins also have many binding partners (Joberty *et al*., 2001; Surka *et al*., 2002; Singh *et al*., 2024). The fact that septins can polymerize into heteromeric high-order structures and bind to numerous binding partners makes them susceptible to perturbation by epitope tags, which are indispensable and widely used to study essentially all proteins including septins.

The fission yeast *Schizosaccharomyces pombe* is an attractive, genetically tractable model organism that has been widely utilized to understand fundamental cellular processes (Balasubramanian *et al*., 2004; Pollard and Wu, 2010; Hoffman *et al*., 2015; Fantes and Hoffman, 2016). *S. pombe* has seven septins (Spn1-7) (Longtine *et al*., 1996; Onishi *et al*., 2010). Spn5-7 only function in sporulation during meiosis, together with Spn2, they form a scaffold, bind to PI(4)P lipid, and guide expansion of the plasma membrane to envelope the forming spores (Onishi *et al*., 2010). Spn1, Spn2, Spn3, and Spn4 are expressed during vegetative growth and localize to the division site as double rings at the rim of the division plane (Longtine *et al*., 1996; Onishi *et al*., 2010). Spn1 and Spn4 are more important for cell division than Spn2 or Spn3, as demonstrated by the deleterious effects of deletion of these septins. Without either Spn1 or Spn4 (in *spn111* or *spn411*), no other septins can localize to the division site (An *et al*., 2004). Spn1-4 are important for septum formation and daughter-cell separation by interacting with the exocyst complex to regulate secretion of the glucanases (Martin-Cuadrado *et al*., 2005; Singh *et al*., 2024). Septin localization at the division site in *S. pombe* is regulated by the anillin Mid2 during cytokinesis (Longtine *et al*., 1996; Onishi *et al*., 2010). However, there are conflicting reports on how septins function and where they localize during interphase. Earlier studies have shown no specific localization of septins on the plasma membrane during interphase (Longtine *et al*., 1996; Onishi *et al*., 2010), while several recent studies have shown that Spn1 and Spn4 localize as puncta or short linear structures all over the plasma membrane, serving as diffusion barriers for proteins such as the NDR kinase Sid2, active Cdc42 GTPase, and Rho GAPs (Zheng *et al*., 2022; McDuffie *et al*., 2024; Zheng *et al*., 2024). It has been proposed that septin puncta or linear filaments on the plasma membrane inhibit the diffusion of Sid2, an NDR-family kinase in the Separation Initiation Network (SIN) pathway, and prevent Sid2’s premature accumulation at the division site (Zheng *et al*., 2018). Additionally, it has been proposed that septins help restrict active Cdc42 to cell tips during interphase for polarized growth (Zheng *et al*., 2022; McDuffie *et al*., 2024).

In this study, we evaluated the localization and function of Spn1 and Spn4 in *S. pombe* when attached to different epitope tags. By analyzing septin localization, cell morphology, cytokinesis/septation defects, cell integrity, and genetic interactions between tagged septins and mutants of the arrestin *art111*, anillin *mid211*, and cold-sensitive *cdc4*-*s16*, we show that Spn1 and Spn4 tagged with mEGFP and mYFP (monomeric YFP with A206K) are more functional and localize to the division site as expected. In contrast, Spn1 and Spn4 tagged with 3HA, tdTomato, and some other red fluorescent tags are less functional. These tags compromise septin localization on the plasma membrane and lead to partial or complete loss of septin function. Our results reemphasize the need to rigorously test the functionality of septins and other proteins with epitope tags. Caution is warranted when interpreting septin function and localization because septin polymers are susceptible to various perturbations.

## RESULTS

### Septins exhibit different localization depending on the epitope tag used

To assess conflicting reports about septin localization, we directly compared the localization of the two most important vegetative septins in *S. pombe* (Spn1 and Spn4) using fluorescence confocal microscopy. Spn1 and Spn4, whose deletions have the strongest cell-division defects (An *et al*., 2004), were each tagged with different fluorescent tags at the C-terminus and expressed under their native promoters at their endogenous chromosomal loci. In the majority of interphase cells expressing Spn1-mYFP or Spn1-mEGFP, Spn1 exhibited diffuse cytoplasmic signals, without any distinct localization on the plasma membrane (Figure 1, A and B). Occasionally, however, Spn1 formed a few stochastic puncta on the plasma membrane. In cells with a partial or complete septum, Spn1 localized to non-contractile rings at the division site (Figure 1, A and B), which could be resolved as double rings at higher spatial resolution (imaging without binning). Spn1 appeared at the division site just before septum appearance as a fuzzy band (Figure 1B, arrowhead), which quickly resolved into septin rings, as the fluorescence intensity in the cytoplasm decreased dramatically (Figure 1, A and B). During daughter-cell separation after septum maturation, Spn1 quickly spread to the new cell ends from the double rings, and then gradually disappeared (Figure 1, A and B). These observations are consistent with previous publications (An *et al*., 2004). In contrast, Spn1-tdTomato localized all over the plasma membrane as puncta or short linear structures in cells at all stages of the cell cycle (Figure 1C). In cells with a forming or closed septum, Spn1-tdTomato localized to the rings at the division site but remained in the structures on the plasma membrane outside the division plane, although with reduced fluorescence intensity and/or incidence (Figure 1C). These observations are also consistent with some previous publications (Zheng *et al*., 2018; Zheng *et al*., 2022; McDuffie *et al*., 2024).

**FIGURE 1:**
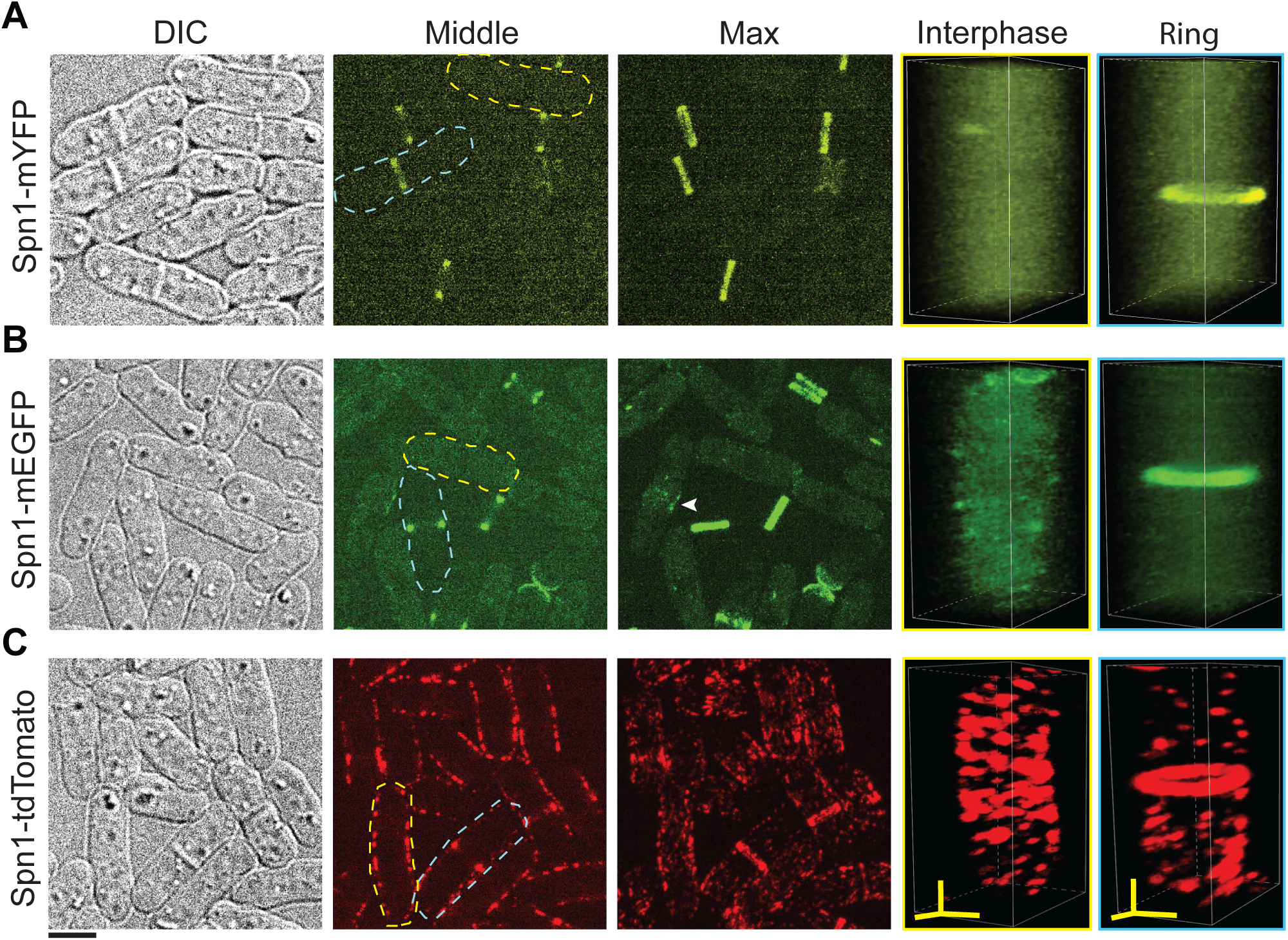
Apparent Spn1 localizations depend on the fluorescent tags. Cell morphology and Spn1 localization in strains expressing Spn1-mYFP (A), Spn1-mEGFP (B), and Spn1-tdTomato (C). Differential Interference Contrast (DIC), fluorescence images at the middle focal plane and the maximal intensity projection of 25 slices with 0.3 μm spacing are shown. 3D volumetric projection of representative interphase and dividing cells (Ring) are on the right. Scale bar, 5 μm. Axial bars, 2 μm.

Tagged Spn4 behaved nearly identically to Spn1 (Figure 2). The localization of Spn4-mYFP and Spn4-GFP(S65T) resembled that of Spn1-mYFP and Spn1-mEGFP, respectively (Figure 2, A and B). Cells expressing Spn4-GFP(S65T) had some puncta on the plasma membrane, which may be due to the fact GFP(S65T) forms a weak dimer (Caviston *et al*., 2003; Oltrogge *et al*., 2014). However, Spn4-GFP(S65T) cells had no obvious septin bars or spirals, which were observed in cells expressing Spn1-YFP but not Spn1-mYFP (Davidson *et al*., 2016). Spn4-tdTomato localized to puncta and short linear filaments all over the plasma membrane, much more than Spn4-GFP(S65T), even during cytokinesis, similarly to Spn1-tdTomato (Figure 2, B and C).

**FIGURE 2:**
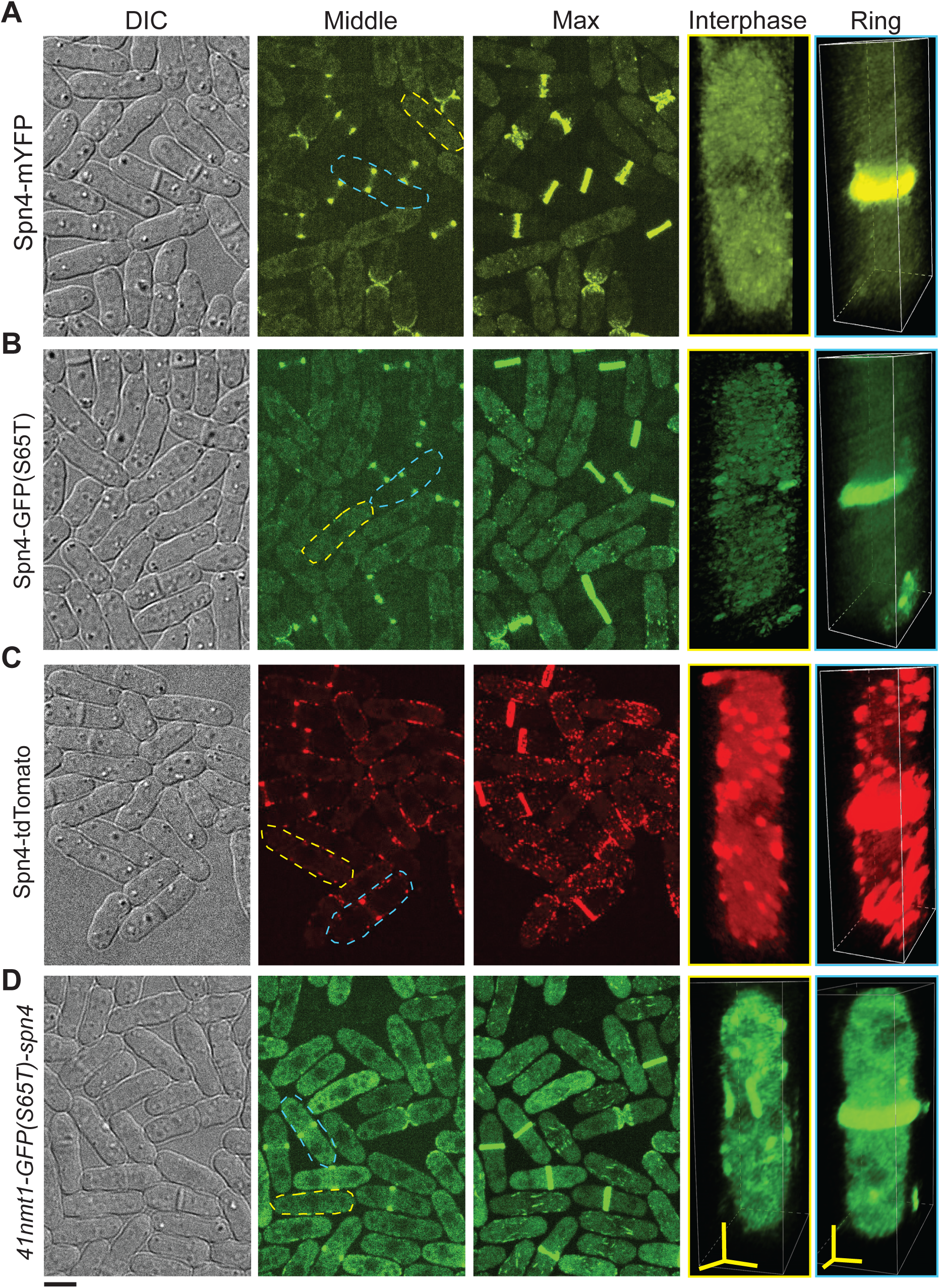
Apparent Spn4 localizations depend on the fluorescent tags. Cell morphology and Spn4 localization in strains expressing Spn4-mYFP (A), Spn4-GFP(S65T) (B), Spn4-tdTomato (C), and GFP(S65T)-Spn4 under the *41nmt1* promoter in YE5S medium (D). DIC, fluorescence images at the middle focal plane and the maximal intensity projection of 25 slices with 0.3 μm spacing are shown. 3D volumetric projection of representative interphase and dividing cells (Ring) are on the right. Scale bar, 5 μm. Axial bars, 2 μm.

To assess if septins exhibit similar localization when tagged at the N-terminus, we used the *S. pombe* strains with integrated *nmt1-GFP(S65T)-spn4* constructs with different promoter strengths. The *nmt1* promoters are regulated by thiamine (Basi *et al*., 1993; Bähler *et al*., 1998). GFP(S65T)-Spn4 expressed at different levels localized to both the division site as rings and short filaments/bundles on the plasma membrane, though the number of filaments appeared less than that of Spn4-tdTomato puncta (Figure 2D and Supplemental Figure S1). The filaments/bundles were mostly along the cell long axis (Figure 2D and Supplemental Figure S1). Moreover, after the septin rings formed, the short filaments almost disappeared from the plasma membrane outside the division site (Figure 2D and Supplemental Figure S1), also behaved differently from Spn4-tdTomato (Figure 2C). These conflicting data on the localizations of Spn1 and Spn4 prompted questions as to which septin localization accurately represents physiological conditions and which is the artifact of epitope tagging.

The mutants of *spn111* and *spn411* exhibited delayed daughter-cell separation and increased septation index (>63%) and cell lysis in a small fraction (2 to 3.5%) of cells (An *et al*., 2004). To test which tagged septins are more functional, we quantified the septation index and cell lysis of cells expressing different fluorescent tags (Table 1). Surprisingly, no significant difference was observed in septation index and cell lysis in cells expressing Spn1 or Spn4 tagged with mEGFP, mYFP, or tdTomato compared to wild type (WT) cells despite their different localizations (Table 1). Thus, it was uncertain which of the septin localizations that we observed so far is real or an artifact.

**Table 1:** Septation index and cell lysis of cells expressing tagged septins in the arrestin *art1^+^* or *art111*.^a^.

| Strain | Mean number of cells scored (n = 3 or 4) | % of cells with the indicated number of septa<sup>b,c</sup> (mean ± SD) |  | % of lysed or shrunken cells (mean ± SD) |
| --- | --- | --- | --- | --- |
|  |  | 1 | ≥2 |  |
| Wild type (WT) | 595 | 15.9 ± 0.5 | 0.00 ± 0.00 | 0.5 ± 0.4 |
| *art1Δ* | 548 | 13.2 ± 1.8 | 0.14 ± 0.24 | 15.9 ± 0.8 |
| *spn1-3HA* | 569 | 63.5 ± 3.3<sup>***</sup> | 1.46 ± 0.27 | 2.3 ± 2.0 |
| *spn1-mEGFP* | 634 | 13.8 ± 4.7 | 0.00 ± 0.00 | 0.4 ± 0.1 |
| *art1Δ spn1-mEGFP* | 656 | 13.0 ± 4.9 | 0.00 ± 0.00 | 17.3 ± 5.1 |
| *spn1-tdTomato* | 595 | 15.7 ± 2.1 | 0.00 ± 0.00 | 0.7 ± 0.2 |
| *spn1-linker-tdTomato* | 565 | 14.0 ± 0.6 | 0.00 ± 0.00 | 0.9 ± 0.5 |
| *art1Δ spn1-tdTomato* | 558 | 17.7 ± 0.9<sup>*</sup> | 0.15 ± 0.17 | 39.2 ± 5.3<sup>†††</sup> |
| *art1Δ spn1-linker-tdTomato* | 523 | 16.2 ± 1.2 | 0.00 ± 0.00 | 18.9 ± 4.0 |
| *spn1-mScarlet* | 640 | 20.0 ± 2.7<sup>*</sup> | 0.00 ± 0.00 | 0.8 ± 0.6 |
| *art1Δ spn1-mScarlet* | 636 | 16.9 ± 7.2 | 0.00 ± 0.00 | 21.9 ± 2.4<sup>†</sup> |
| *spn1Δ* | 521 | 62.9 ± 7.2<sup>**</sup> | 2.31 ± 0.69 | 2.0 ± 2.3 |
| *art1Δ spn1Δ* | 638 | 54.5 ± 15.8<sup>*</sup> | 7.64 ± 3.89 | 46.2 ± 2.3<sup>†††</sup> |
| *spn4-3HA* | 790 | 52.0 ± 8.6<sup>**</sup> | 1.42 ± 0.47 | 1.3 ± 0.8 |
| *spn4-mYFP* | 576 | 13.9 ± 2.8 | 0.00 ± 0.00 | 0.4 ± 0.4 |
| *art1Δ spn4-mYFP* | 597 | 12.7 ± 4.2 | 0.00 ± 0.00 | 21.9 ± 5.3 |
| *spn4-tdTomato* | 508 | 16.0 ± 2.4 | 0.00 ± 0.00 | 0.7 ± 0.7 |
| *art1Δ spn4-tdTomato* | 645 | 15.3 ± 2.4 | 0.00 ± 0.00 | 29.3 ± 3.3<sup>††</sup> |
| *spn4-mScarlet* | 546 | 32.7 ± 3.2<sup>***</sup> | 0.13 ± 0.11 | 1.4 ± 0.4 |
| *art1Δ spn4-mScarlet* | 644 | 30.8 ± 5.4<sup>**</sup> | 0.39 ± 0.40 | 24.5 ± 0.4<sup>†††</sup> |
| *spn4Δ* | 562 | 65.8 ± 5.6<sup>***</sup> | 2.76 ± 1.87 | 3.5 ± 3.5 |
| *art1Δ spn4Δ* | 524 | 57.5 ± 4.5<sup>**</sup> | 10.33 ± 5.90 | 39.5 ± 8.4<sup>††</sup> |
| *spn4-mYFP spn1-tdTomato* | 586 | 13.9 ± 2.4 | 0.00 ± 0.00 | 0.7 ± 0.2 |
| *spn4-tdTomato spn1-mEGFP* | 693 | 16.0 ± 2.8 | 0.04 ± 0.06 | 0.2 ± 0.3 |
| *art1Δ spn4-tdTomato spn1-mEGFP* | 550 | 14.9 ± 4.6 | 0.00 ± 0.00 | 27.1 ± 0.8<sup>†††</sup> |
| *art1Δ spn1-mYFP<sup>d</sup>* | 583 | 18.6 ± 1.4<sup>*</sup> | 0.00 ± 0.00 | 14.5 ± 1.9 |
| *art1Δ spn1-YFP<sup>d</sup>* | 578 | 17.8 ± 1.3 | 0.00 ± 0.00 | 15.2 ± 5.2 |

<sup>a</sup>Cells were grown in log phase (OD<sub>595</sub> < 0.5) in YE5S liquid medium for ~36 h at 25°C before imaging.
<sup>b</sup>The lysed or shrunken cells were excluded from the septation index calculation.
<sup>c</sup>Septa were counted using DIC images, which could lead to an underestimate because the emerging septum may not be easily visible.
<sup>d</sup>The *p* values of the septation index for *art1Δ spn1-mYFP* = 0.036 and *art1Δ spn1-YFP* = 0.082.
<sup>\*</sup>The septation index (total septa/total cells) is significantly greater than WT cells in two tailed Student's *t*-test. <sup>\*</sup>*p* < 0.05, <sup>\*\*</sup>*p* < 0.01, <sup>\*\*\*</sup>*p* < 0.001.
<sup>†</sup>The percentage of lysed or shrunken cells is significantly greater than *art1Δ* cells in *t*-test. <sup>†</sup>*p* < 0.05, <sup>††</sup>*p* < 0.01, <sup>†††</sup>*p* < 0.001.

### A flexible linker partially restores Spn1-tdTomato localization to the division site but a small 3HA tag almost abolishes septin localizations and functions

To assess if a flexible linker between a septin protein and the epitope tag would alter the localizations of septin fusion proteins to make them more consistent, we inserted a long flexible linker between Spn1 and mYFP, mEGFP, or tdTomato. The linker <u>ILGAPSGGGATAGAGGAGGPAGLI</u> was previously used to construct functional EB1-GFP, which does not interfere with its binding to microtubules (Sandblad *et al*., 2006). Cells expressing Spn1-linker-mYFP or Spn1-linker-mEGFP under their native promoter at endogenous chromosomal locus exhibited similar division-site localization as Spn1-mYFP and Spn1-mEGFP without the linker but had slightly more puncta or linear structures on the plasma membrane, especially for Spn1-linker-mEGFP (Supplemental Figure S2, A and B). In contrast, the addition of the linker to Spn1-tdTomato resulted in decreased localization to puncta or short linear structures on the plasma membrane and increased diffuse cytoplasmic signals during interphase and brighter Spn1 rings at the division site (Supplemental Figure S2C; Supplemental Movie 1). Additionally, the kinetics of septin puncta during cell division were altered in Spn1-linker-tdTomato cells. When Spn1-linker-tdTomato concentrated at the division site to form rings, puncta nearly completely faded from the plasma membrane outside the division site, which is significantly different from Spn1-tdTomato (Supplemental Figure S2, C and D; Supplemental Movie 1). During daughter-cell separation, Spn1-linker-tdTomato exhibited increased diffusion into the cytoplasm from the cell ends than Spn1-tdTomato cells (Movie 1). Thus, the addition of a flexible linker does modify the localization of tdTomato tagged septins.

The tdTomato tag (54.2 kDa) is twice as big as GFP or YFP (27.0 kDa). To ask whether epitope tag size influences septin localization and function, we attached the much smaller 3HA tag (4.5 kDa) to the C-terminus of both Spn1 and Spn4 (Bähler *et al*., 1998). Surprisingly, cells expressing Spn1-3HA or Spn4-3HA phenotypically resembled *spn111* and *spn411* cells, exhibiting a much higher septation index than *spn211* or *spn311* cells (Figure 3A and Table 1). Approximately 16% of WT *S. pombe* cells had a visible single septum, and no cells had multiple septa in an asynchronous population under our growth conditions (Table 1). In cells with 3HA tagged septins, the percentage of cells with a septum increased to ∼64% and ∼52% for Spn1-3HA and Spn4-3HA, respectively, and ∼1.5% of cells contained two or more septa (Table 1 and Figure 3A). These defects were worse when cells were grown on plates, indicating that some septated cells were separated by shaking in the liquid cultures. In addition, more cells had a curved morphology, opposed to the typical straight rod shape. This resulted in cells overlapping on slides during microscopy and led to an underestimate of cellular defects in quantification (Figure 3A and Table 1). We conclude that *spn1-3HA* and *spn4-3HA* cells are defective in completing cell division, and that epitope tag size is not solely responsible for the defects in septin localization and function.

**FIGURE 3:**
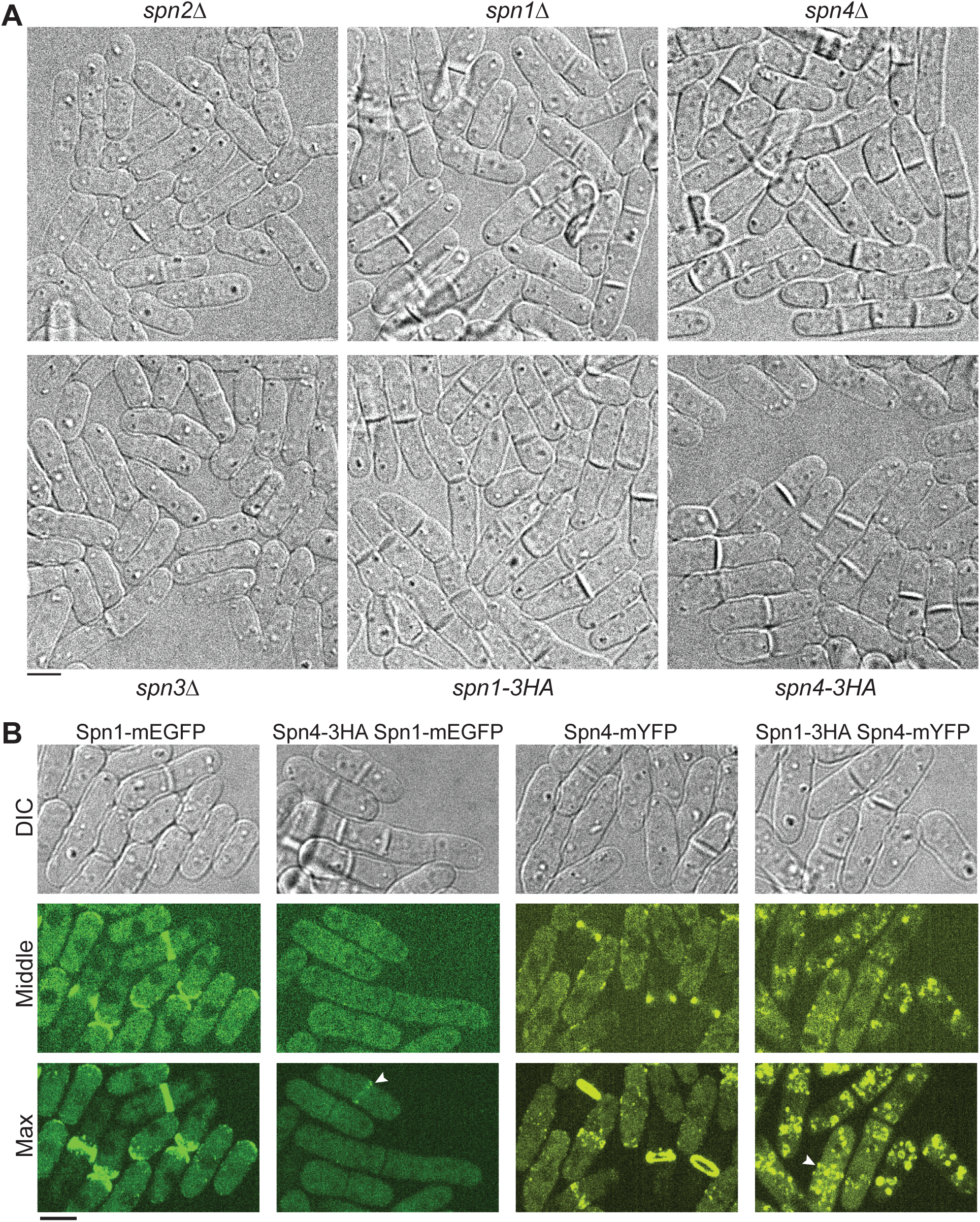
Small 3HA tag abolishes septin functions and localizations. (A) DIC images showing the morphology of *spn211, spn111, spn411, spn311, spn1-3HA*, and *spn4-3HA* cells grown at 25°C. (B) The 3HA tag essentially abolishes septin localizations to the division site and the plasma membrane. DIC, middle focal plane, and maximal intensity projections of 25 slices with 0.3 μm spacing of cells expressing Spn1-mEGFP, Spn4-3HA Spn1-mEGFP, Spn4-mYFP, or Spn1-3HA Spn4-mYFP are shown. Arrowheads mark the faint septin rings. Scale bars, 5 μm.

The delay in daughter-cell separation and the high septation index of Spn1-3HA or Spn4-3HA expressing cells led us to hypothesize that septins do not localize properly to the plasma membrane in these cells. Indeed, fewer than 2% of cells exhibited ring localization of the septin Spn1-mEGFP or Spn4-mYFP in cells expressing Spn4-3HA or Spn1-3HA, respectively (Figure 3B). The residual ring was much fainter and incomplete compared to WT cells (Figure 3B; marked with arrowheads). No other signals were detected on the plasma membrane, and Spn4-mYFP formed aggregates in the cytoplasm (Figure 3B). Thus, the 3HA tag essentially abolishes septin functions and their localization to the plasma membrane in fission yeast.

### Tagged septins affect the localization of other septins

Because 3HA compromises septin localization (Figure 3B), we next tested whether septins colocalize in different structures on the plasma membrane, and what the effect of using various epitope tags (tdTomato, mEGFP, GFP[S65T], and mYFP) would have on (co)-localization since septins form hetero-oligomers. We found that the septins Spn1 and Spn4 colocalized in all structures on the plasma membrane, including puncta, linear structures, and rings regardless of the epitope tag combinations used (Figure 4, A-C). Additionally, we determined that tagged Spn1 is the dominant septin; other septins followed the localization pattern of Spn1 under all conditions during interphase (Figure 4, A-C). Strikingly, Spn4-tdTomato puncta and linear structures on the plasma membrane during interphase or outside the division site during cell division mostly disappeared in cells co-expressing Spn1-mEGFP (Figure 4C), suggesting that Spn1-mEGFP affects the localization of Spn4-tdTomato differently than untagged Spn1, and therefore, Spn1-mEGFP is not 100% functional. However, cell morphology of all the three strains had no obvious differences compared to WT (Figure 4). Moreover, the septation index and percentage of cell lysis in cells expressing Spn4-mYFP Spn1-tdTomato or Spn4-tdTomato Spn1-mEGFP were not significantly different from WT (Table 1). These results highlight the complex nature of studying septin localizations/interactions and the need for rigorous tests to determine the localizations and functionalities of tagged septins.

**FIGURE 4:**
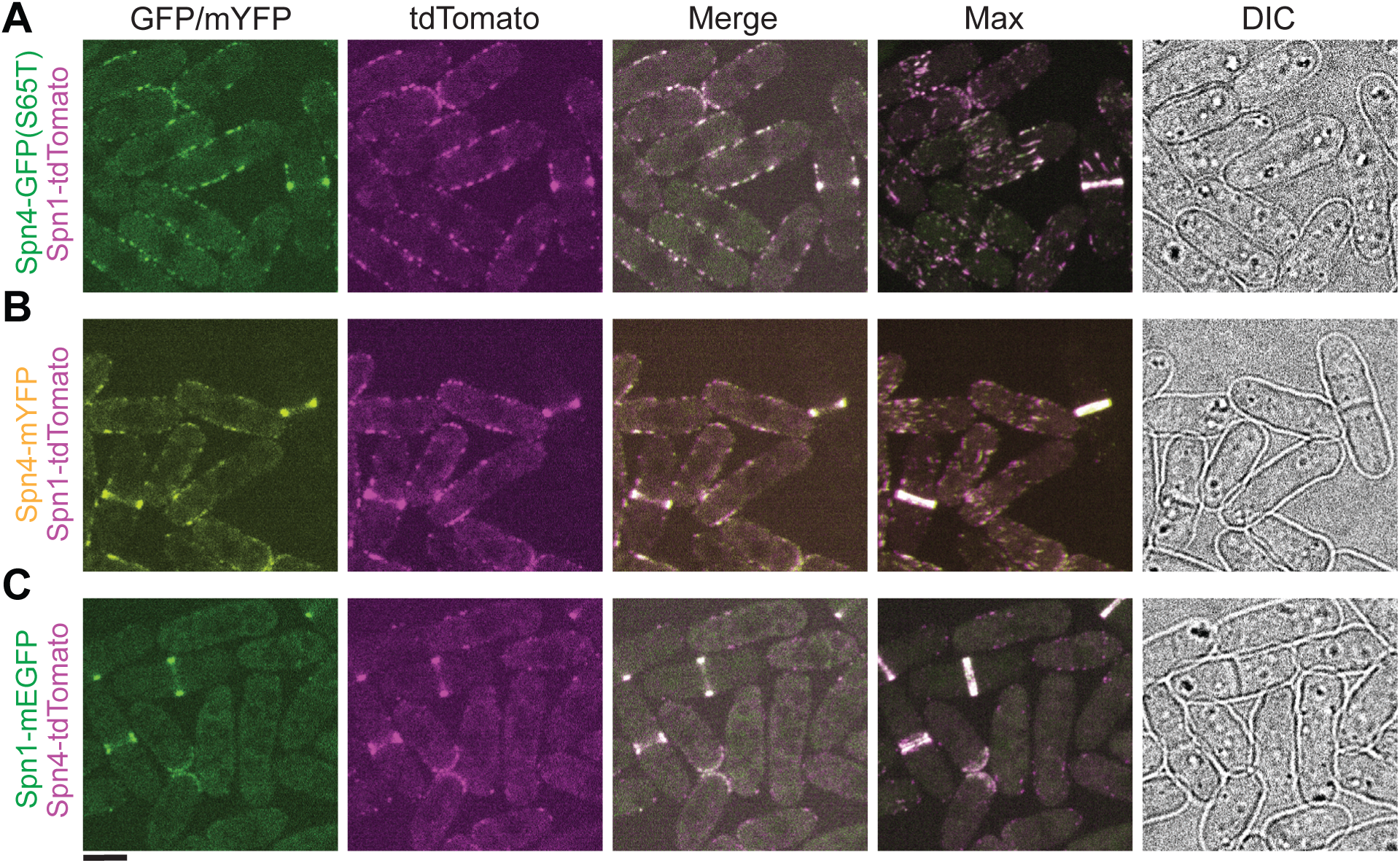
Tagged septins can affect each other’s localizations. Colocalization of Spn4-GFP(S65T) and Spn1-tdTomato (A), Spn4-mYFP and Spn1-tdTomato (B), and Spn1-mEGFP and Spn4-tdTomato (C). From left to right: middle focal plane of GFP/mYFP, tdTomato, and merged channels; the maximal intensity projection of 25 slices spaced at 0.3 μm for the merged channels; and DIC. Scale bar, 5 μm.

### Septin localization to plasma membrane puncta or linear structures is independent of the anillin Mid2

The anillin Mid2 colocalizes with septins at the division site and is known to be important for septin assembly and organization (Berlin *et al*., 2003; Tasto *et al*., 2003; Wu *et al*., 2010; Arbizzani *et al*., 2022). We tested if the formation of septin puncta outside the division site on the plasma membrane depends on Mid2. Since *spn1* and *mid2* genes are tightly linked (<104 kb apart) on the same chromosome, we could not obtain the *spn1-tdTomato mid211* strain so we tested Spn4 only. In *mid2^+^* cells, Spn4-mYFP localized to the division site as rings during cytokinesis, with very faint signals in the cytoplasm and on the plasma membrane (Figure 5, A and C). During interphase, Spn4-mYFP displayed diffuse cytoplasmic signals as well as localization in several puncta on the plasma membrane (Figure 5, A and C). This observation is consistent with Figure 2A. In *mid211* cells, Spn4-mYFP remained diffuse in the cytoplasm during interphase. However, during cytokinesis, Spn4-mYFP intensity in the rings decreased, and the signals in the cytoplasm and on the plasma membrane increased (Figure 5, A and C), consistent with previous reports (Berlin *et al*., 2003; Tasto *et al*., 2003; Wu *et al*., 2010; Arbizzani *et al*., 2022). The division site signal of Spn4-tdTomato was also significantly decreased during cytokinesis in *mid211* cells. However, Spn4-tdTomato still formed puncta and linear structures on the plasma membrane with high intensity during interphase and outside the division site during cytokinesis in *mid211* cells as in *mid2^+^* cells (Figure 5, B and C). Consistently, Mid2 only localized to the rings during cell division, not to the puncta or linear structures outside the division site or during interphase (Figure 5D). Thus, we concluded that the cortical localization of Spn4-tdTomato outside the division site, unlike the septin localization in the rings, is Mid2 independent. Our data presented so far give no convincing inferences as to which septin localizations are native and which are artifacts. Thus, more sensitive tests are required to discern the functionality of tagged septins.

**FIGURE 5:**
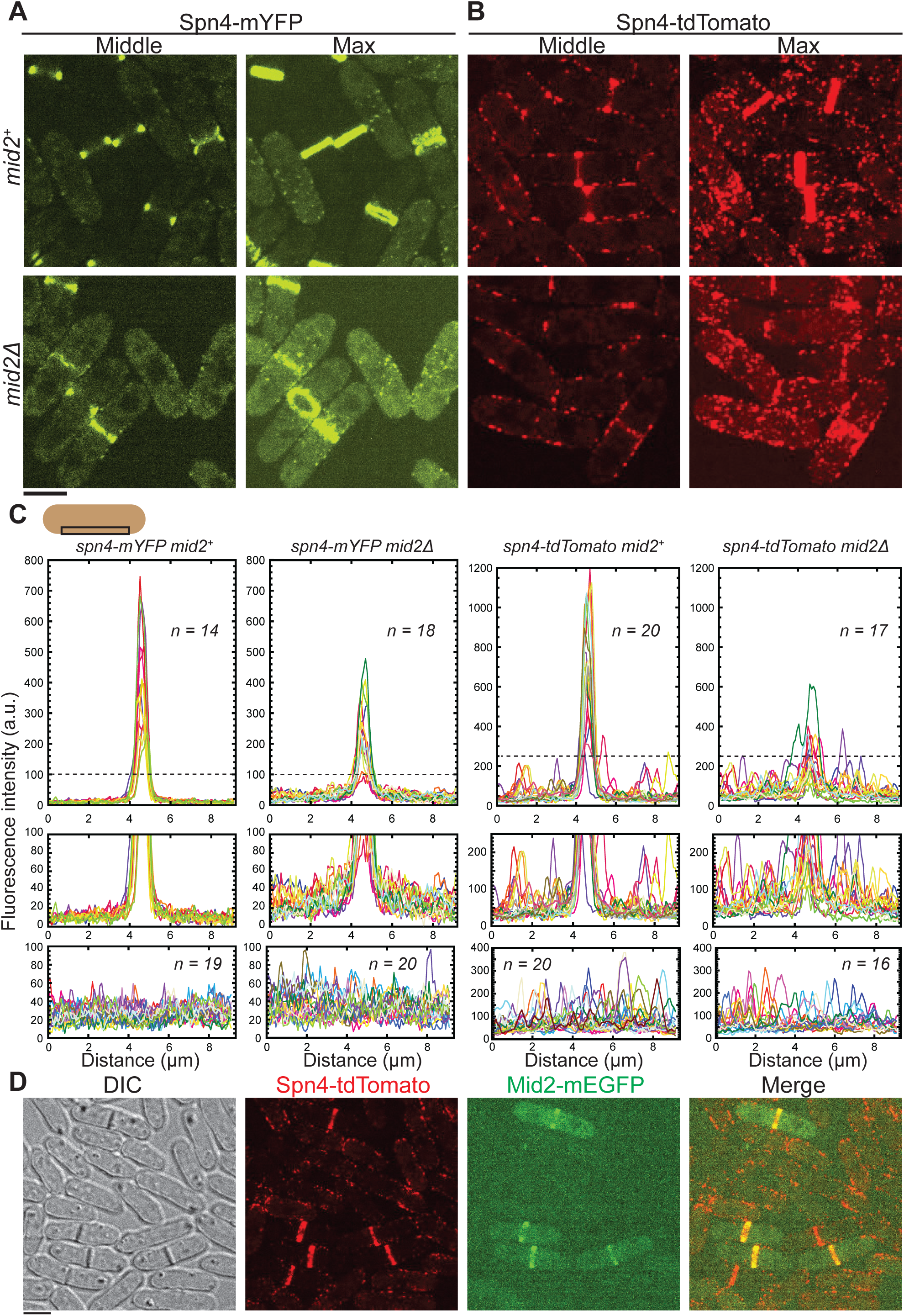
Localization of Spn4 to the division site but not to the interphase membrane puncta or linear structures depends on the anillin Mid2. Localization of Spn4-mYFP (A) and Spn4-tdTomato (B) in *mid2^+^* (top) and *mid211* (bottom) cells. Images of the middle focal plane and maximal intensity projection of 25 slices with 0.3 μm spacing are shown. (C) Plots of fluorescence intensity on the plasma membrane measured using the middle focal plane along the cell-long axis (9.2 μm long centered at the division site within a width of 0.92 μm as shown by the schematic) in cells with the septin rings (top and middle) and in interphase cells (bottom). The dashed line in the top graph marks the portion of the graph enlarged in the middle graphs to view the signal outside the division site with higher resolution. (D) Mid2-mEGFP colocalizes with Spn4-tdTomato at the division-site rings but not at the puncta or linear filaments outside the division site. DIC, and maximal intensity projection of 25 slices with 0.3 μm spacing (for the fluorescence images) are shown. Scale bars, 5 μm.

### Genetic interactions between tagged septins and the arrestin *art1****11*** or cold-sensitive *cdc4-s16* reveal that tdTomato tagged septins are less functional

The cell morphology of all the strains expressing tagged septins examined so far (except 3HA) resembled WT (Figures 1–5). None of the cells expressing Spn1-mEGFP, Spn1-mYFP, Spn1-tdTomato, Spn4-mYFP, or Spn4-tdTomato showed a significant difference in either cell lysis or septation index from WT (Table 1 and Figure 6, A-C, upper panels). This indicates that quantification of these two parameters alone is not sufficient to distinguish which fluorescently tagged septins retain their native localization and normal function. Additionally, the growth assay using plates with Phloxin B— which stains lysed or dead cells red because they cannot pump out the red dye— failed to distinguish tagged septin strains at temperatures ranging from 20 to 36°C. Thus, it was necessary to use more sensitive genetic interactions to test the functionality of the tagged septins.

**FIGURE 6:**
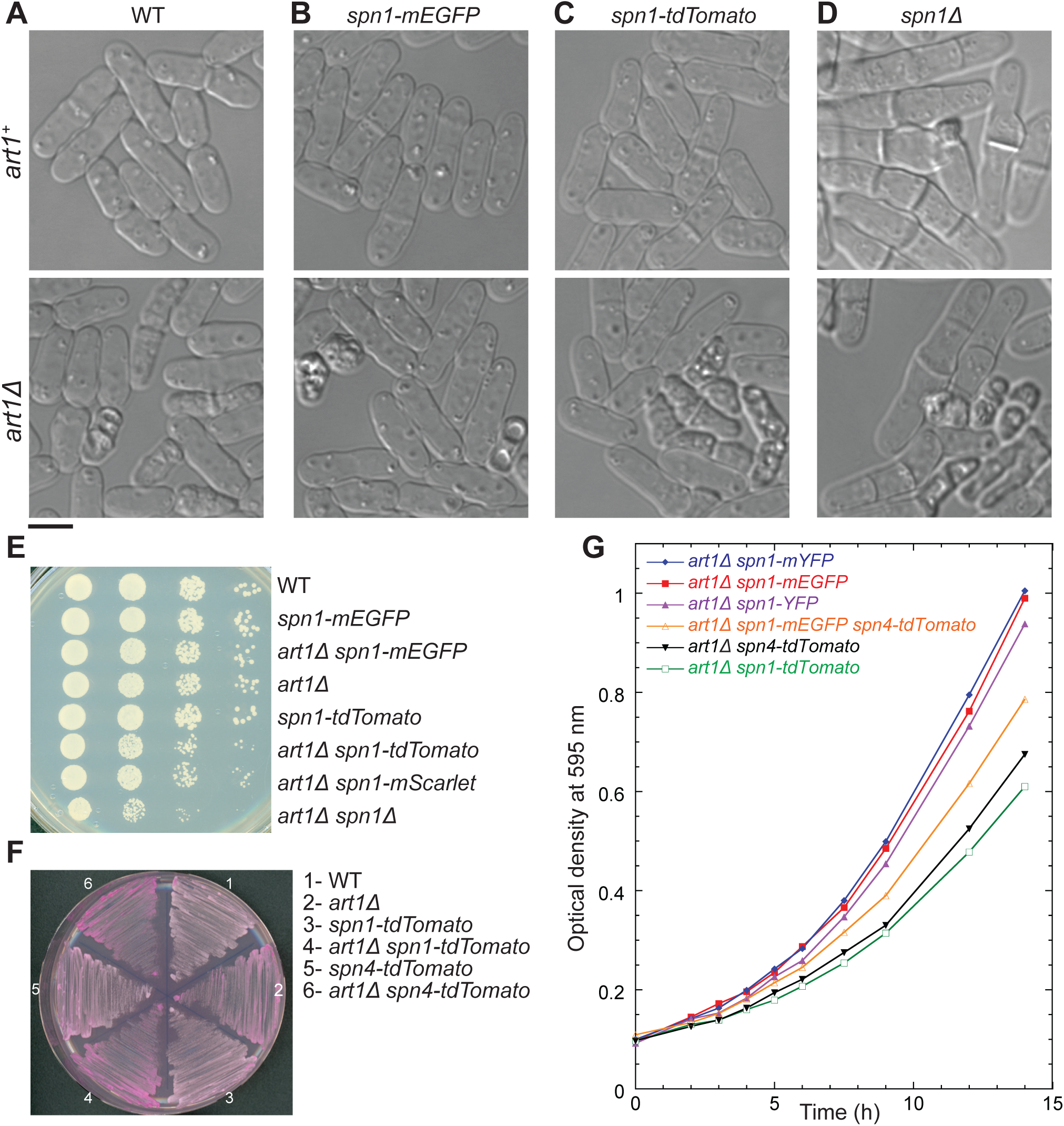
Synthetic genetic interactions between the arrestin *art111* and tagged septins or *spn111* and cell growth of the indicated strains. (A-D) Cell morphology under DIC of WT (A), *spn1-mEGFP* (B), *spn1-tdTomato* (C), and *spn111* (D) cells in WT *art1*^+^ (top) or *art111* (bottom) cells. Scale bar, 5 μm. (E) Serial dilutions (10x) to test growth of the eight indicated strains. Cells were grown exponentially in YE5S for ∼36 h before spotting onto YE5S plates. The plates were grown for ∼48 h at 36°C. (F) Synthetic genetic interactions between *art111* and *spn1-tdTomato* or *spn4-tdTomato.* Cells were grown at 25°C for 48 h on YE5S plate with Phloxin B, which stains unhealthy, lysed, or dead cells red. Darker red color indicates more cell lysis. (G) Growth curves of the indicated strains by measuring the optical density at 595 nm across a 14 h period. Strains were all grown in YE5S liquid media at 25°C.

The arrestin-like protein Art1 and myosin light chain Cdc4 function in parallel pathways with septins to regulate cytokinesis and cell integrity (Wu *et al*., 2010; Davidson *et al*., 2015). The double mutants of septins with either a*rt111* or *cdc4-s16* lead to significant cell lysis during daughter-cell separation and/or severely defective septum formation (Wu *et al*., 2010). To investigate the subtle effects of tagged septins on cell health, we crossed fluorescently tagged septin strains with *art111* or cold-sensitive *cdc4-s16* (Wu *et al*., 2010). We hypothesized that strains expressing more functional septins would display a similar phenotype as the *art1Δ* or *cdc4-s16* mutant. In contrast, cells expressing less functional septin fusion proteins would have more severe synthetic phenotypes. We found that *art111 spn1-mEGFP* and *art1Δ spn4-mYFP* cells exhibited similar phenotypes as *art111* in terms of percentage of cell lysis and septation index (Table 1 and Figure 6, A and B). In contrast, ∼39% *art1Δ spn1-tdTomato* and 29% *art1Δ spn4-tdTomato* cells were lysed, which is significantly different from *art1Δ* (∼16%) cells (Table 1). *art1Δ spn1-tdTomato* cells resembled the cell lysis rates of *art1Δ spn1Δ* and *art1Δ spn4Δ* cells, which had 46% and 40% of cells lysed, respectively (Table 1 and Figure 6, C and D). However, the septation index was much higher in *art1Δ spn1Δ* and *art1Δ spn4Δ* cells (Table 1).

Serial dilutions and growth on plates with Phloxin B confirmed our results (Figure 6, E and F). In the serial dilutions, *art1Δ spn1-tdTomato* cells grew less than both *art111 spn1-mEGFP* and *art111* based on the number and size of colonies at 36°C (Figure 6E), 32°C, and 28°C. Additionally, *art1Δ spn1-tdTomato* and *art1Δ spn4-tdTomato* cells had more cell lysis than either single mutant, which was evident from the darker red color (meaning the cells cannot pump out the red dye) on Phloxin B plate grown at 25°C (Figure 6F). Moreover, growth curves measured using liquid cultures grown at 25°C confirmed that *art111 spn1-mEGFP* and *art111 spn1-mYFP* cells grew much faster than *art1Δ spn1-tdTomato* and *art1Δ spn4-tdTomato* cells (Figure 6G). Thus, mEGFP and mYFP tagged septins Spn1 and Spn4 are more functional than tdTomato tagged septins in this arrestin Art1 parallel pathway in regulating cell division and cell integrity. In addition, cells expressing Spn1-mScarlet and Spn4-mScarlet also had higher septation index than WT (Table 1), and the localization pattern of Spn1-mCherry was similar to that of Spn1-tdTomato (unpublished data). Collectively, these data suggest that mEGFP and mYFP tagged septins are more functional than septins tagged with tdTomato or other red fluorescent proteins in completing cytokinesis and maintaining cell integrity.

We confirmed that tdTomato tagged septins are less functional than septins tagged with mEGFP and mYFP using the cold-sensitive *cdc4-s16* mutant (Wu *et al*., 2010). *cdc4-s16* cells expressing Spn1-tdTomato or Spn4-tdTomato had more cell lysis, higher septation index, more partial septa, and branched cells than *cdc4-s16* cells with or without Spn1-mEGFP and Spn4-mYFP (Table 2 and Supplemental Figure S3). However, *cdc4-s16* cells expressing either Spn1-mEGFP or Spn4-mYFP were more defective in septation (the septation index is significantly different) and cell integrity than *cdc4-s16* cells alone (Table 2 and Supplemental Figure S3), which suggested that mEGFP and mYFP tagged septins are also not fully functional in this Cdc4 parallel pathway in regulating cytokinesis and septation.

**Table 2:**
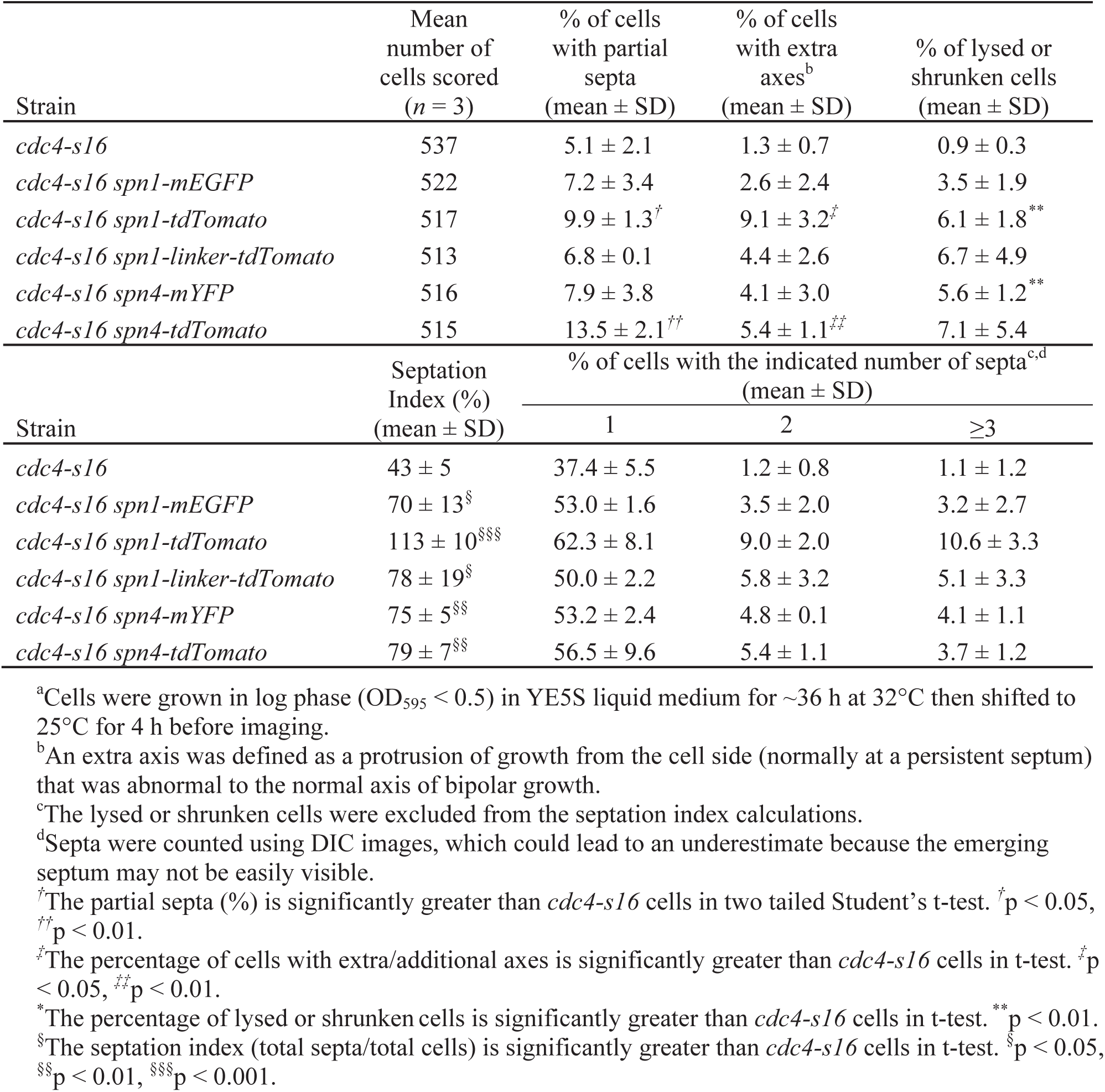
Septum defects and cell lysis in cold-sensitive *cdc4-s16* mutant cells expressing tagged septins.^a^.

| Strain | Mean<br>number of<br>cells scored<br>(*n* = 3) | % of cells<br>with partial<br>septa<br>(mean ± SD) | % of cells<br>with extra<br>axes<sup>b</sup><br>(mean ± SD) | % of lysed or<br>shrunk cells<br>(mean ± SD) |
| --- | --- | --- | --- | --- |
| *cdc4-s16* | 537 | 5.1 ± 2.1 | 1.3 ± 0.7 | 0.9 ± 0.3 |
| *cdc4-s16 spn1-mEGFP* | 522 | 7.2 ± 3.4 | 2.6 ± 2.4 | 3.5 ± 1.9 |
| *cdc4-s16 spn1-tdTomato* | 517 | 9.9 ± 1.3<sup>†</sup> | 9.1 ± 3.2<sup>‡</sup> | 6.1 ± 1.8<sup>**</sup> |
| *cdc4-s16 spn1-linker-tdTomato* | 513 | 6.8 ± 0.1 | 4.4 ± 2.6 | 6.7 ± 4.9 |
| *cdc4-s16 spn4-mYFP* | 516 | 7.9 ± 3.8 | 4.1 ± 3.0 | 5.6 ± 1.2<sup>**</sup> |
| *cdc4-s16 spn4-tdTomato* | 515 | 13.5 ± 2.1<sup>††</sup> | 5.4 ± 1.1<sup>‡‡</sup> | 7.1 ± 5.4 |
| Strain | Septation<br>Index (%)<br>(mean ± SD) | % of cells with the indicated number of septa<sup>c,d</sup><br>(mean ± SD) |  |  |
|  |  | 1 | 2 | ≥3 |
| *cdc4-s16* | 43 ± 5 | 37.4 ± 5.5 | 1.2 ± 0.8 | 1.1 ± 1.2 |
| *cdc4-s16 spn1-mEGFP* | 70 ± 13<sup>§</sup> | 53.0 ± 1.6 | 3.5 ± 2.0 | 3.2 ± 2.7 |
| *cdc4-s16 spn1-tdTomato* | 113 ± 10<sup>§§§</sup> | 62.3 ± 8.1 | 9.0 ± 2.0 | 10.6 ± 3.3 |
| *cdc4-s16 spn1-linker-tdTomato* | 78 ± 19<sup>§</sup> | 50.0 ± 2.2 | 5.8 ± 3.2 | 5.1 ± 3.3 |
| *cdc4-s16 spn4-mYFP* | 75 ± 5<sup>§§</sup> | 53.2 ± 2.4 | 4.8 ± 0.1 | 4.1 ± 1.1 |
| *cdc4-s16 spn4-tdTomato* | 79 ± 7<sup>§§</sup> | 56.5 ± 9.6 | 5.4 ± 1.1 | 3.7 ± 1.2 |

<sup>a</sup>Cells were grown in log phase (OD<sub>595</sub> < 0.5) in YE5S liquid medium for ~36 h at 32°C then shifted to 25°C for 4 h before imaging.
<sup>b</sup>An extra axis was defined as a protrusion of growth from the cell side (normally at a persistent septum) that was abnormal to the normal axis of bipolar growth.
<sup>c</sup>The lysed or shrunk cells were excluded from the septation index calculations.
<sup>d</sup>Septa were counted using DIC images, which could lead to an underestimate because the emerging septum may not be easily visible.
<sup>†</sup>The partial septa (%) is significantly greater than *cdc4-s16* cells in two tailed Student's t-test. <sup>†</sup>*p* < 0.05, <sup>††</sup>*p* < 0.01.
<sup>‡</sup>The percentage of cells with extra/additional axes is significantly greater than *cdc4-s16* cells in t-test. <sup>‡</sup>*p* < 0.05, <sup>‡‡</sup>*p* < 0.01.
<sup>\*</sup>The percentage of lysed or shrunk cells is significantly greater than *cdc4-s16* cells in t-test. <sup>\*\*</sup>*p* < 0.01.
<sup>§</sup>The septation index (total septa/total cells) is significantly greater than *cdc4-s16* cells in t-test. <sup>§</sup>*p* < 0.05, <sup>§§</sup>*p* < 0.01, <sup>§§§</sup>*p* < 0.001.

Lastly, we used the sensitized *art111* and/or *cdc4-s16* genetic backgrounds to resolve three of the unanswered questions about septin localization in fission yeast. First, we tested if Spn1-linker-tdTomato is more functional than Spn1-tdTomato because the localization of Spn1-linker-tdTomato is more similar to Spn1-mEGFP/mYFP than Spn1-tdTomato (Figure 1 and Supplemental Figure S2). Judging by septation index, percentage cell lysis, septum and cell morphology in *art111* and *cdc4-s16* mutants (Tables 1 and 2), Spn1-linker-tdTomato was more functional than Spn1-tdTomato, though not fully functional. Thus, the flexible linker does improve Spn1 localization and function. Second, in Figure 4C, we showed that Spn1-mEGFP partially prevents Spn4-tdTomato from localizing to the puncta and short linear filaments on the plasma membrane outside the division site. We tested if cells expressing both Spn1-mEGFP and Spn4-tdTomato are healthier than expressing Spn4-tdTomato alone in *art111* background. Although *art111* did not obviously affect the localizations of Spn4-tdTomato or Spn4-tdTomato Spn1-mEGFP (compare Supplemental Figure S4A and Figure 4C), and there was no significant difference in septation index and cell lysis between the two strains (Table 1). Interestingly, though, *art111 spn4-tdTomato spn1-mEGFP* cells grew slightly faster than *art111 spn4-tdTomato* cells (Figure 6G). Third, we reported that Spn1-YFP but not Spn1-mYFP forms septin bars and spirals in the cytoplasm, and that these structures persist even in cells with septin rings during cell division (Davidson *et al*., 2016). We investigated which Spn1 is more functional using the *art111* background. The localizations of Spn1-YFP and Spn1-mYFP were the same in *art1^+^* (Davidson *et al*., 2016) and *art111* cells (Supplemental Figure S4, B and C). Thus, Art1 does not affect the localization or mislocalization of tagged Spn1. Surprisingly, *art111 spn1-mYFP* and *art111 spn1-YFP* cells grew at the same rate and had very similar septation index and percentage of cell lysis (Figure 6G and Table 1). The similarity between these two strains demonstrates that not all aspects of septin localization and function can be distinguished by functional complementation or by using a sensitized genetic background. The tagged septins may not have a phenotype even if they are mislocalized or form an artifact in the cells.

### Localization of the NDR kinase Sid2 to the division site is delayed in septin mutants

Models have been proposed that septin puncta and linear structures on the plasma membrane in cells expressing Spn1-tdTomato or Spn4-tdTomato serve as a diffusion barrier that affects the localization and function of the NDR kinase Sid2, the Rho GAPs Rga6 and Rga4, and active Cdc42 (Zheng *et al*., 2018; Zheng *et al*., 2022; McDuffie *et al*., 2024). Because we have found that tdTomato tagged septins are less functional compared to mEGFP or mYFP tagged septins, we reexamined the relationship between Sid2 and septins.

Sid2 is the most downstream kinase in the SIN (septation initiation network) pathway, whose homologs function in the equivalent MEN pathway in budding yeast and the Hippo pathway in animal cells including humans (McCollum and Gould, 2001; Yoshida *et al*., 2002; Hergovich and Hemmings, 2012; Heallen *et al*., 2013; Hsu and Weiss, 2013; Foltman and Sanchez-Diaz, 2017; Sinha *et al*., 2022). Sid2 localizes to the spindle pole body (SPB) throughout the cell cycle and to the division site during cytokinesis (Sparks *et al*., 1999). A previously proposed model suggests that the septin-tdTomato puncta on the plasma membrane inhibit Sid2’s localization to the division site and help maintain SIN signaling once the septin rings are formed (Zheng *et al*., 2018). In *spn111* cells, Sid2 appearance at the division site was reportedly significantly advanced, and Sid2 duration at the cleavage furrow was significantly shortened compared to WT (Zheng *et al*., 2018). Using time-lapse confocal microscopy, we found that both Spn1-mEGFP and Spn4-mYFP appeared in the rings at the division site ∼6 minutes earlier than Sid2-mEGFP in WT cells (Figure 7, A-C). In *spn111* and *spn411* cells, Sid2 arrived at the division site at 32.2 ± 3.8 and 31.5 ± 3.8 min after SPB separation, respectively, which was delayed compared to 29.7 ± 3.0 in WT cells (Figure 7, D and E; p = 0.002 and 0.028). Additionally, we observed that Sid2 persisted at the division site similarly in WT and *spn111* cells, whereas Sid2 disappeared earlier in *spn411* cells (Figure 7F). In WT and *spn1Δ* cells, Sid2 departed from the division site at ∼38 min and ∼37 min after its appearance, respectively, while in *spn4Δ* cells Sid2 stayed for ∼33 min (Fig 7F). Together, the findings that septins appear at the division site earlier than Sid2 in WT cells and that Sid2 appearance is delayed in septin deletion mutants do not support the model that septins serve as a diffusion barrier to prevent Sid2 localization to the division site.

**FIGURE 7:**
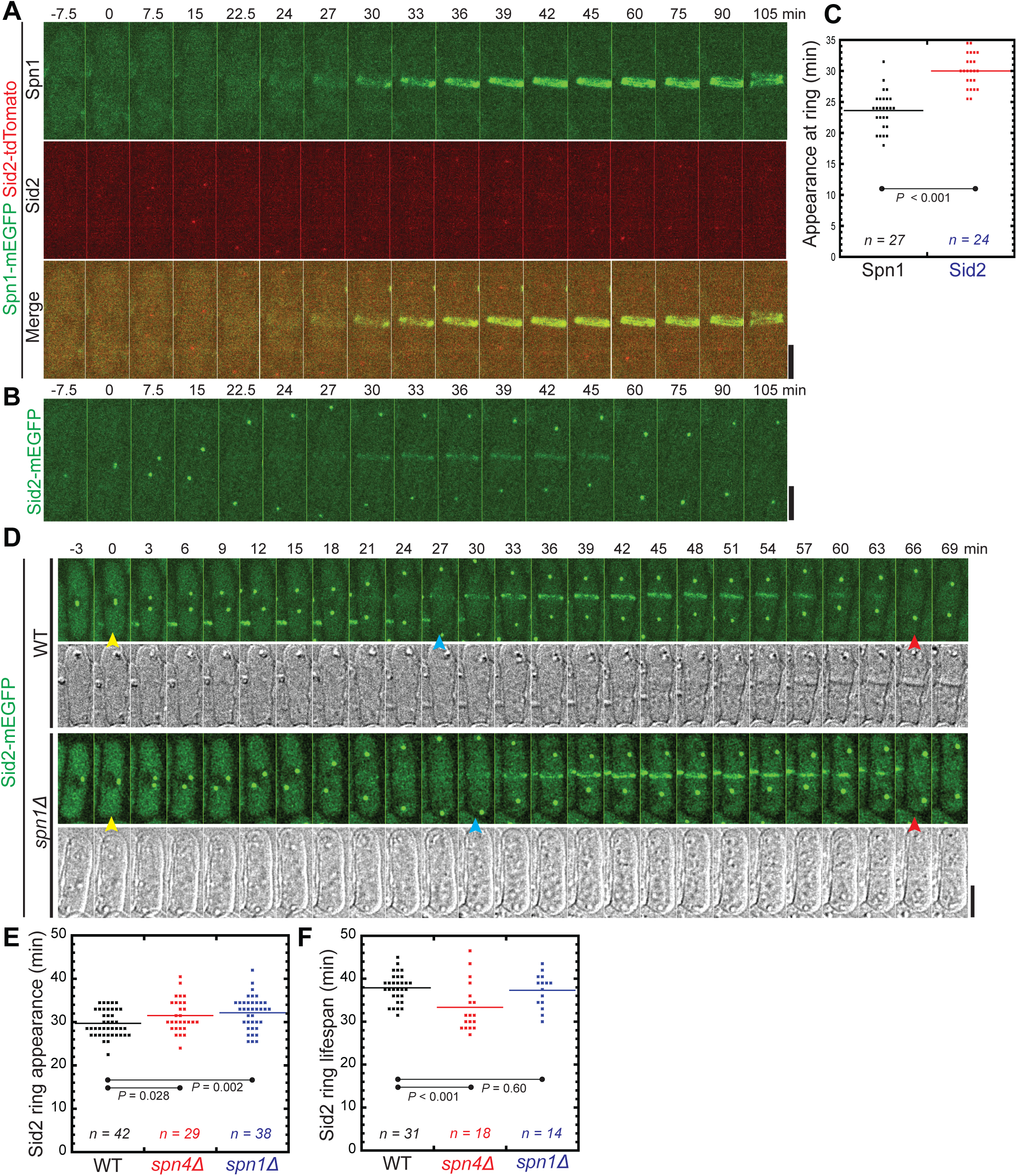
Sid2 localization at the division site in WT and septin mutants. (A-C) Time courses (A and B) and quantification (C) of Spn1 and Sid2 appearance at the division site. Time courses of maximal intensity projection of 9 slices with 0.9 μm spacing for Spn1 (A) and Sid2 (B). Cells expressing both Spn1-mEGFP and Sid2-tdTomato or Sid2-mEGFP alone were mixed at ∼1:1 ratio and imaged on the same slide to make sure the imaging conditions are identical. SPB separation as marked by Sid2 is defined as time 0. Because Sid2-tdTomato (as SPB marker for Spn1 quantification) signal is weak intentionally due to using very low laser power, Sid2-mEGFP was used to quantify Sid2 appearance in (C). (D-F) Sid2 appearance and duration at the division site in WT and septin mutants. (D) Time courses of maximal intensity projection of 13 slices with 0.5 μm spacing of Sid2-mEGFP in WT and *spn1Δ* cells. Yellow arrows: SPB separation at the onset of mitosis; Blue arrows: Sid2 arrival at division site; and Red arrows: Sid2 disappearance from the division site. Scale bars, 5 μm. (E and F) Quantification of (E) Sid2-mEGFP ring appearance at the division site (SPB separation defined as time 0) and (F) the lifespan of Sid2-mEGFP (from Sid2 ring appearance to its departure from the division site) in WT, *spn411*, and *spn111* cells. P values were calculated by two-tailed Student’s t-test.

### Epitope tags are predicted to affect the octameric structure of septin filaments

We utilized AlphaFold3, which is capable of accurately modeling protein folding and interactions within the biomolecular space through a single unified deep-learning framework (Jumper *et al*., 2021; Abramson *et al*., 2024), to predict the octameric septin filamentous structure and probe the interactions between septin subunits when Spn1 and/or Spn4 are epitope-tagged. Our predictions of the non-epitope tagged septin octamer, or WT, are consistent with previously reported septin structures (Cavini *et al*., 2021; Singh *et al*., 2024). The septin subunits are ordered Spn3-Spn4-Spn1-Spn2-Spn2-Spn1-Spn4-Spn3 with an iPTM score of 0.47 and a pTM score of 0.52 (Figure 8A), which is consistent with the established organization of the *S. pombe* septin and reminiscent of the budding yeast palindromic septin ordering of Cdc11-Cdc12-Cdc3-Cdc10-Cdc10-Cdc3-Cdc12-Cdc11 (An *et al*., 2004; Versele *et al*., 2004; Bertin *et al*., 2008). Moreover, the interfacial binding motif of the WT octamer appears to follow the canonical pattern of the G-domain (GTP-binding domain) binding between Spn3 and Spn4, N-C domain binding of Spn4 and Spn1, G-domain binding of Spn1 and Spn2, and the N-C domain binding of the two Spn2 at the central core (Figure 8A) (Cavini *et al*., 2021). Therefore, our predicted models are supported by previous experimental findings (Longtine *et al*., 1996; An *et al*., 2004).

**FIGURE 8:**
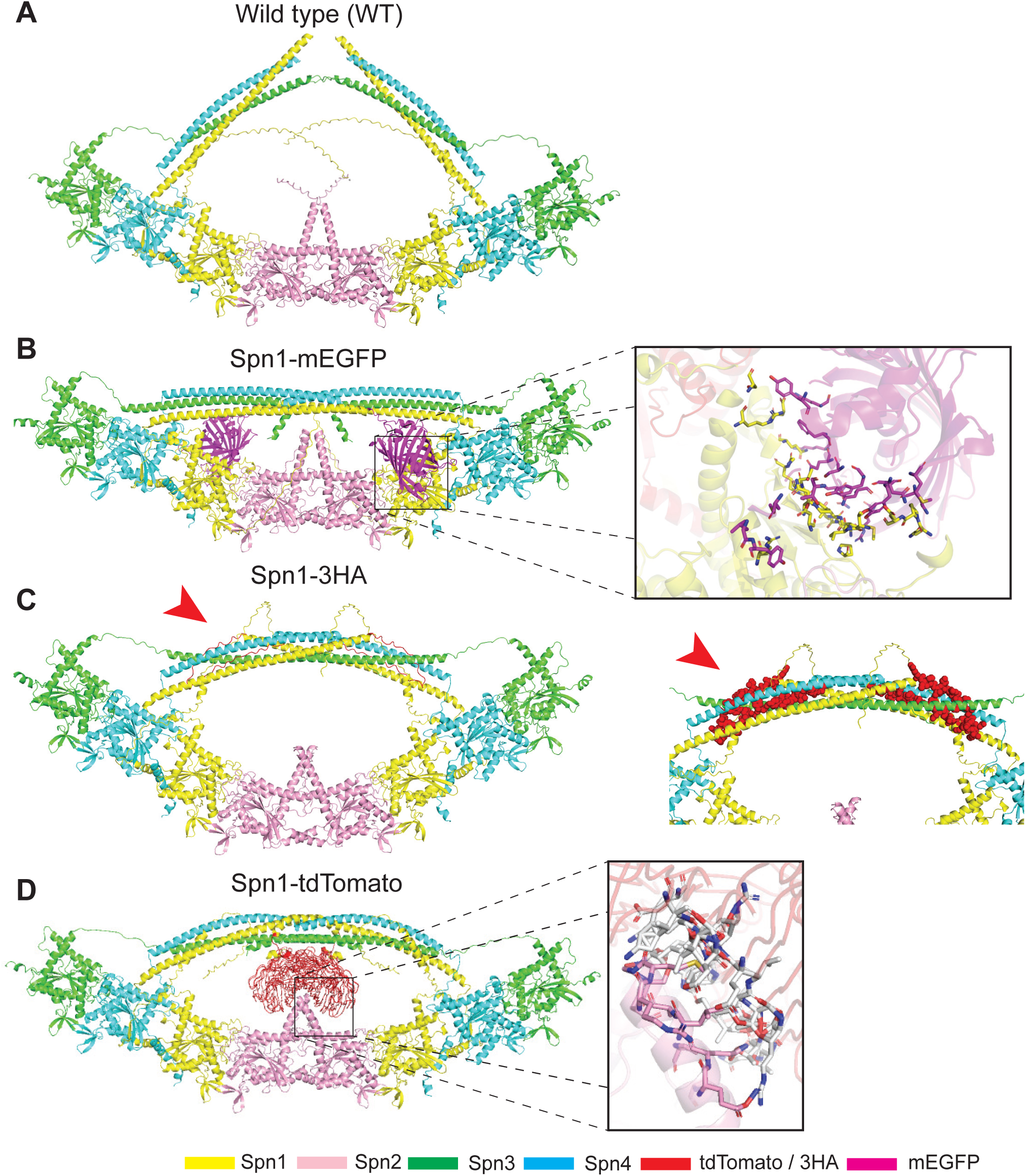
Predicted structures of the WT septin octameric filament and epitope-tagged Spn1 by AlphaFold3. (A-D) Septins Spn1-4 are differentiated by the color as labeled: Spn1, yellow; Spn2, pink; Spn3, green; and Spn4, blue. Two subunits of each septin were used to construct the octameric filament. The tandem-dimer tomato (tdTomato) and 3HA (3X hemagglutinin) are colored in BR9 (red), and the mEGFP subunit is colored in magenta. Red arrowheads indicate the 3HA interaction with the bundled coiled coils. (B and D) Box projections show interacting residues between (B) mEGFP (magenta) and Spn1 (yellow) or (D) tdTomato (white) and Spn2 (magenta) within 3.5Å, indicative of Van der Waals forces represented as sticks. (A) pTM = 0.52. (B) pTM = 0.48. (C) pTM = 0.48. (D) pTM = 0.49.

Epitope tagging of Spn1 changes the structure of the octameric filament. Adding mEGFP to Spn1 does not lead to significant changes in the core octameric filament but results in the bundling of coiled-coil domains of Spn1, Spn3, and Spn4 (Figure 8B). In addition, the model predicts that certain residues in mEGFP and Spn1 would be within 3.5Å from one another, which would be in the range for them to interact via Van der Waals forces (Figure 8B, Box Projection). Interestingly, Spn1-3HA also imparts a slight bending effect on septin filaments in addition to the bundling of the coiled-coil domains (Figure 8C). No significant changes are observed in the octameric filament of our Spn1-tdTomato model, but the software predicts an interaction between tdTomato and the C-terminal domain of the Spn2 core, which also leads to tighter coiled-coil bundling (Figure 8D). AlphaFold3 predicts that >24 residues from the tdTomato tag interact with 11 residues from the C-terminal domain of Spn2 via Van der Waals interactions within a 3.5Å distance (Figure 8D, Box projection).

Our Spn4 models follow a distinct pattern. The tags in both Spn4-mYFP and Spn4-GFP(S65T) are predicted to interact with Spn1 to confer a bending of the octameric filament and bundling of the coiled-coils, which may be responsible for the slight difference observed between the Spn1 and Spn4 localization as well as the larger variation in cell lysis in tagged Spn4 strains (Fig. 1 and 2; Table 1; Supplemental Figure S5, A and B). Prediction of Spn4-3HA shows no change in the filament structure and no tight bundling in which the Spn1, Spn3, and Spn4 coiled-coil domains appear to prefer a “peak” conformation as opposed to its “bowed” Spn1-3HA counterpart (Supplemental Figure S5C). Our Spn4-tdTomato model confers a greater bend in filament structure in comparison to Spn1-tdTomato but shows the same interactions between tdTomato and the Spn2 C-terminal domain (Supplemental Figure S5D). Similar Van der Waals interactions are inferred within a 3.5Å distance between 27 residues from the tdTomato subunit and 12 residues from the C-terminal domain of Spn2 (Supplemental Figure S5D, Box projection).

To further investigate the mechanism underpinning the dominating effect of epitope-tagged Spn1 on the localization of other septins (Figure 4C; Spn1-mEGFP affecting Spn4-tdTomato localization), we predicted the structures for Spn1-mEGFP Spn4-tdTomato and Spn1-tdTomato Spn4-GFP(S65T). Both models appear to confer a significant bend in the octameric filament (Supplemental Figure S6, A and B). When Spn1 is tagged with mEGFP, the allosteric inhibition between tdTomato and mEGFP is significant. tdTomato partially interacts with the Spn2 C-terminal interface to sterically hinder mEGFP from interacting with Spn1 (Supplemental Figure S6A). tdTomato consists of two monomeric subunits that are covalently link together via a 12-residue linker (GHGTGSTGSGSS) to allow some flexibility (Shaner *et al*., 2004). In our predicted model, one monomeric unit of tdTomato interacts with the Spn2 C-terminal interface while the other prevents mEGFP from interacting with Spn1 as previously described (Figure 8B), which results in a separation of the coiled-coil bundle (Supplemental Figure S6A). Thus, the bundling of the coiled-coils is reduced. Conversely, when Spn1 is tagged with tdTomato, we also see allosteric inhibition between the epitope tags, but Spn4-GFP(S65T) is more closely situated on the coiled-coils preventing tdTomato from interacting with the Spn2 C-terminal interface which results in a tetrameric-like formation of the tdTomato monomers and significantly increases coiled-coil bundling (Supplemental Figure S6B). Interestingly, our Spn1-mYFP model predicts a specific interaction between mYFP and the opposing Spn1 protein, which causes an insignificant change in the main octameric filament but a bundling of the coiled-coils (Supplemental Figure S6C). In contrast, weak dimeric Spn1-YFP containing octamer is bent at the Spn2 core, mediated by an apparent dimerization of the two centered YFP subunits, which is situated above the Spn2 core and promotes bundling of the coiled-coils (Supplemental Figure S6D). Collectively, the AlphaFold3 predictions are consistent with our conclusions that no tagged septins are 100% functional, tdTomato is more disruptive to septin structures than mEGFP and mYFP, and monomeric fluorescent proteins are less likely to affect septin structures than weak dimeric forms.

## DISCUSSION

In this study, we investigated the discrepancy over the localization and function of septins in fission yeast (An *et al*., 2004; Zheng *et al*., 2018; Zheng *et al*., 2022; McDuffie *et al*., 2024). Though all tagged septin proteins were expressed at their native chromosomal loci and under the control of their native promoters (except *nmt1-GFP[S65T]-spn4*), we found that Spn1 and Spn4 tagged with tdTomato or mScarlet were less functional than their mEGFP or mYFP tagged counterparts. Moreover, even mEGFP or mYFP tagged Spn1 and Spn4 are not fully (or 100%) functional. In addition, we found that the small tag, 3HA, nearly abolishes septin localization and function as cells expressing Spn1-3HA or Spn4-3HA resemble septin deletion mutants, and septins cannot efficiently localize to the plasma membrane. Our data suggest that previously reported septin puncta and linear filaments/structures on the plasma membrane or septin bars in the cytoplasm are likely caused by non-physiological septin fusion proteins with decreased functionality, which interfere with other septins’ localization and function. It is not likely that these structures serve as diffusion barrier for proteins involved in cytokinesis and cell polarity as they are too labile.

### Epitope tags may induce septin mislocalization and dysfunction

By assessing septin fusion proteins in live fission yeast cells, we highlight the necessity of using caution when interpreting localization and function data of septins. We find that several different epitope tags can perturb both the function and localization of septins. Several lines of evidence indicate that mEGFP and mYFP-tagged septins are more functional and localized normally to the double rings at the division site, whereas tdTomato and 3HA-tagged septins suffer from partial to complete loss of function. First, cells expressing Spn1 or Spn4 tagged with mEGFP or mYFP resemble WT cells in morphology, septation index, cell integrity, growth at different temperatures, and have no synthetic genetic interactions with the arrestin *art111* (Figures 1, 2, 6 and Table 1). These tagged septins only localize to the division site as non-constricted rings from late anaphase until daughter-cell separation. This localization pattern is consistent with previous results from the Pringle laboratory, which used “affinity-purified antibodies that specifically recognized one of the *S. pombe* septins Spn1, Spn2, or Spn3, found by immunofluorescence microscopy of fixed cells that all three septins localized to the septal region at the time of cell division, whereas no localized staining of any of these three proteins was observed during other stages of the cell cycle” (Longtine *et al.,* 1996; J.R. Pringle, personal communication). Second, cells expressing 3HA-tagged Spn1 and Spn4 resemble *spn1Δ* or *spn4Δ* in percent of cells with defects in septum formation, delay in daughter-cell separation, and cell lysis (Figure 3 and Table 1). Although 3HA is a small tag relative to the other fluorescent proteins used, it almost disrupts septin localization and function (Figure 3B). Third, cells expressing Spn1 or Spn4 with the tdTomato tag resemble WT cells in their morphology and septation index. However, in double mutants with the arrestin *art111*, these cells have 247% or 84% increase in cell lysis, respectively, compared to *art111* cells (Table 1) while mEGFP or mYFP tagged Spn1 or Spn4 have no significant synthetic genetic interactions with *art111*. When combined with the cold-sensitive *cdc4-s16*, the tdTomato tagged Spn1 or Spn4 are more defective in septum formation, cytokinesis, and cell lysis than mEGFP and mYFP tagged septins (Table 2 and Supplemental Figure S3). Because Spn1-tdTomato and Spn4-tdTomato spread all over the plasma membrane, even during cytokinesis, the levels of these septins at the division site are much lower than cells with mEGFP or mYFP tagged septins (Figures 1, 2, and 5). It is likely that the reduced septin level at the division site cannot sustain septin function in cytokinesis without Art1. Thus, tdTomato and 3HA tags lead to partial or complete loss of septin function. The genetic interactions with *cdc4-s16* also revealed that Spn1-mEGFP and Spn4-mYFP are not 100% functional since their double mutants with *cdc4-s16* are more defective in septation and cell lysis than *cdc4-s16* (Table 2 and Supplemental Figure S3). Although the localizations of Spn1-mEGFP and Spn4-mYFP are consistent with immunostaining using septin-specific antibodies (Longtine *et al.,* 1996; J.R. Pringle, personal communication), the fluorescent tags may still affect septin binding or dynamics on the plasma membrane, or affect septin assembly or structure, which is suggested by our AlphaFold3 modeling (Figure 8 and Supplemental Figures S5 and S6).

The septin puncta or short linear filaments/structures across the plasma membrane outside the division site in cells expressing tdTomato tagged septins have been proposed to serve as diffusion barriers (An *et al*., 2004; Zheng *et al*., 2018; Zheng *et al*., 2022; McDuffie *et al*., 2024). However, our data are not consistent with that model for several reasons. First, Sid2 appears at the division site later in septin deletion mutants than in WT cells (Figure 7, D and E). Additionally, septins appear at the division site several minutes earlier than Sid2 in WT cells (Figure 7, A-C), which is not consistent with the model that septins inhibit Sid2 localization to the division site. However, since Sid2 is less abundant than septins at the division site (Figure 7, A and B), we cannot rule out that Sid2 appears at the division site several minutes earlier than what we were able to detect. We find this scenario less likely, however, since Sid2-tdTomato signal appears at the division site at a similar time as Sid2-mEGFP, despite its weak signal. We think that the septins are necessary for the arrival of Sid2 to the division site instead of preventing its arrival. Therefore, it appears that septins may serve as a scaffold at the division site for recruiting Sid2. Second, cells expressing Spn1-mEGFP Spn4-tdTomato and Spn4-mYFP Spn1-tdTomato have very similar morphology, septation index, and very low cell-lysis percentage (Table 1) despite Spn1 and Spn4 localizations being dramatically different outside the division site in these cells (Figure 4), which suggests that the septin puncta and short linear filaments do not play important roles. Third, the anillin Mid2 is crucial in septin localization and organization at the division site during cytokinesis (Berlin *et al*., 2003; Tasto *et al*., 2003; Wu *et al*., 2010; Arbizzani *et al*., 2022); however, Spn4-tdTomato localization on the plasma membrane outside the division site does not depend on Mid2 (Figure 6). Thus, it is more likely that septin-tdTomato puncta and linear structures during interphase or outside the equator during cell division have no significant functional role. Consistently, a flexible linker can partially restore Spn1-tdTomato localization and function, making it function more similarly to Spn1-mEGFP in *art111* cells, especially during cell division (Supplemental Figure S2; Tables 1 and 2). Septin mislocalization uniformly to all over the plasma membrane has been observed in budding yeast upon overexpression of certain septin subunits (Benson and McMurray, 2023). Although the sensitized *art111* genetic background cannot help distinguish if the constructs are functional or not (Supplemental Figure S4, B and C; and Table 1), other septin structures that we observed, such as septin bars or spirals in cells expressing Spn1-YFP, or short filaments in cells expressing GFP(S65T)-Spn4, may also have no physiological roles. Collectively, septin hetero-oligomers are susceptible to perturbation by epitope tags, which leads to the formation of various cellular structures on the plasma membrane and in the cytoplasm. But it seems that not all of the septin structures that we and others have observed are physiologically relevant. However, septin structures, such as septin patches on the plasma membrane outside the division site, have been observed in budding yeast using super-resolution microscopy or during autophagy (Barve *et al*., 2018; Perucho-Jaimes *et al*., 2024; El Alaoui *et al*., 2025); therefore, it remains to be investigated if any septin structures besides the double rings at the division site have any functions in fission yeast.

### Why various epitope tags have drastically different effects on septin localization and function?

It is likely that different epitope tags interfere with or affect the interactions between septin subunits, septins with septin-binding proteins, and septins with the plasma membrane differently. Septins form palindromic, oligomeric structures when they bind to one another at both the NC interface and the G interface (Sirajuddin *et al*., 2007). It is likely that 3HA and tdTomato tags, and to a lesser extent mEGFP and mYFP, interfere with the NC interface since all the tags are added C-terminally. It is possible that epitope tags, such as tdTomato, trigger a conformational change in septin subunits that lead to a loss in septin curvature or lipid preference. Consistently, various tags do induce conformational changes in septin octamer structures, especially the bundling or packing of the coiled-coil motifs. Intriguingly, attachment of tdTomato, but not other tags, to Spn1 or Spn4 causes interaction with Spn2, which induces tighter bundling and packing of the coiled-coil motifs (Figure 8; Supplemental Figures S5 and S6). It is clear that tdTomato compromises the membrane specificity of both Spn1 and Spn4. It is also possible that epitope tags induce the same structural changes that mimic a post-translational modification. These possibilities may explain the mixed results that we observed when expressing two differently tagged septins in the same cell (Figure 4). It has also been shown that Spn1-YFP, but not Spn1-mYFP (monomeric YFP with A206K), causes the formation of straight, long septin bars or spirals in fission yeast (Davidson *et al*., 2016), and these structures are Art1 independent (Supplemental Figure S4, B and C). This corroborates the idea that septin localization and assembly are susceptible to epitope tagging.

Septins bind to the plasma membrane by interacting with anionic lipids like PI(4,5)P2 and phosphatidylserine and then polymerizing into filaments (Zhang *et al*., 1999; Bridges *et al*., 2014). The theoretical pI of 3HA is 3.17, and its aliphatic index is 39.00 (Ikai, 1980), which is more acidic than mEGFP, mYFP, and tdTomato (pI: 5.79, 6.06, and 6.31 respectively). Thus, 3HA likely prevents septins from binding to the negatively charged plasma membrane or disrupts septin filament assembly. Indeed, AlphaFold3 modeling suggests 3HA causes bigger conformational changes on septin filaments (Figure 8 and Supplemental Figure S5). Septins also preferentially recognize and interact with the plasma membrane with micron-scale curvature (Bridges *et al*., 2016). In rod-shaped fission yeast cells, the membrane curvatures at the cleavage furrow, cell sides, and cell tips are quite different. Interestingly, short septin filaments in cells overexpressing GFP(S65T)-Spn1 are mostly parallel to cell long axis where membrane curvature is the least, which suggests that these septin structures have lost curvature preference. The localization and function of septins are also regulated by phosphorylation and ubiquitination (Johnson and Blobel, 1999; Vargas-Muñiz *et al*., 2016). Future studies are required to investigate the effects of epitope tags in septin membrane specificity, as different epitope tags clearly affect determinants of septin localization differently.

### Functionality tests of tagged septins in other systems

Septins play essential roles in cytokinesis, cell polarization, sporulation, and many other cellular processes across diverse cell types by forming numerous higher order structures (Garcia *et al*., 2011). In cell types other than fungi, it is technically challenging to test the functionality of widely used epitope tagged septins, even when these septins are expressed at their endogenous levels. Moreover, as shown in our study of Spn1-YFP, some tagged septins may not exhibit a unique phenotype, even in a sensitized genetic background. Most known functionality tests have been done in budding yeast or other fungi by rescue mutant phenotype or by observing cell morphology (Chang, 1999; Johnson and Blobel, 1999). The budding yeast has at least two advantages in evaluating the functionality of tagged septins compared to fission yeast and most other cell types. First, most septins are essential in budding yeast, and multiple temperature-sensitive alleles are available. Non-functionally tagged septins do not survive or would show obvious phenotypes. Second, the pioneering septin localization studies were done using septin-specific antibodies before the discovery of GFP and other fluorescent proteins. Thus, non-functional septin fusion proteins that cause mislocalization or artifact are less likely to appear in literature. When GFP fusions of Cdc3, Cdc10, Cdc11, and Cdc12 were first utilized in budding yeast, there were varying levels of assessments of their functionality. Studies utilizing Cdc10-GFP and Cdc3-GFP made no direct statements about the functionality of these fusion proteins (Cid *et al*., 1998; Moffat and Andrews, 2004). A study utilizing all four fusion proteins claimed that all four were fully functional (Takahashi *et al*., 1999). However, Chang et al. reported that Cdc12-GFP does not completely restore *cdc12* loss-of-function mutant phenotype (Chang, 1999). Moreover, even one of the most functional tagged septins in budding yeast, GFP-Cdc3, is not 100% functional (Caviston *et al*., 2003).

The functionalities of tagged septins have been evaluated in many other fungal model systems. In *M. oryzae*, the tagged septin was examined for functionality by deletion complementation (Dagdas *et al*., 2012). In *A. nidulans*, tagged septins were assessed by an array of impressive characterizations such as the number of nuclei, septa, branches, germ tubes, growth states, and conidiophore morphology (Hernández-Rodríguez *et al*., 2012). However, as shown in our study, Spn1-tdTomato would have passed most of those tests. It was only in a stressed state (some mutant backgrounds) under which the functional defects was revealed. Indeed, it has been shown that tagged septins are not fully functional under challenged conditions via chitinase and bipolar assessment assays in *U. maydis* (Zander *et al*., 2016).

In summary, our study highlights that even for septins expressed under their native promoters, as the sole copy in a cell, 1) the localization of the same septin can be dramatically different with different tags; 2) it is likely many of the tagged septins used in this or similar studies are not 100% functional; 3) some of the septin structures or localizations observed are likely not physiological relevant and have no cellular functions; 4) monomeric fluorescent proteins, such as mEGFP/mYFP (A206K), cause less perturbations than their weak dimer counterparts; and 5) rigorous functionality tests are necessary, especially for cytoskeletal polymers.

Our study reinforces a vital need for rigorous functionality tests of epitope tagged proteins and caution in interpreting localization data when a rigorous test is not feasible. It has been previously shown that fluorescent tags can alter the localization and binding interactions of proteins (Snapp, 2009; Gao *et al*., 2022; Fatti *et al*., 2025). In the context of condensate formation, tagging Dhh1, an mRNA degradation factor enriched in P-bodies, with GFP or mCherry2 inhibits its normal condensation ability (Fatti *et al*., 2025). Moreover, epitope tagging of G-actin monomers interferes with its incorporation into linear actin filaments nucleated by formins (Doyle and Botstein, 1996; Wu and Pollard, 2005; Chen *et al*., 2012). Many cytokinesis proteins are not functional when tagged at their COOH termini (Wu *et al*., 2003). Because epitope tagged proteins are widely used and indispensable for cellular and biochemical studies of septins and essentially all other proteins, caution must be used when interpreting protein localization, interactions, and functions of fusion proteins, especially for proteins that polymerize into heteromeric high-order structures, such as septins.

## MATERIALS AND METHODS

### Yeast strains and genetic methods

The fission yeast *S. pombe* strains used in this study are listed in Supplemental Table S1. The yeast strains were woken up from -80℃ stocks onto YE5S plates and grown at 25℃ (or 32°C for cold-sensitive *cdc4-s16* strains) for 2 to 3 days before experiments. Gene targeting was performed using the standard PCR-based homologous recombination method (Bähler *et al*., 1998). All the epitope tagged genes (except *nmt1*-*GFP[S65T]-spn4*) are controlled by their native promoters and the *ADH1* terminator and integrated at their normal chromosomal loci. The *GFP[S65T]* gene controlled by the *nmt1* promoter of different strengths was inserted before the start codon of *spn4* (Bähler *et al*., 1998). To construct double mutants, crosses and tetrad dissection were carried out according to the standard genetic method (Moreno *et al*., 1991). Briefly, freshly growing cells of opposite mating type (*h^+^* or *h^-^*) were mixed in sterile water on an SPA5S plate and incubated at 25℃ (30°C for *cdc4-s16* strains) for ∼48 h. The cross was streaked atop a YE5S plate for tetrad dissection using a Nikon 50i dissection microscope, and the plate was incubated at 25°C (or 32°C for *cdc4-s16* strains) for 4 to 7 days. The colonies of sufficient sizes were picked up using sterile toothpicks onto a new YE5S plate and grown for one or two additional days. The plates were then replica plated onto YE5S selection plates with antibiotics to identify the double mutants.

The flexible linker <u>ILGAPSGGGATAGAGGAGGPAGLI</u> was successfully used to construct the functional EB1-GFP fusion protein (Sandblad *et al*., 2006). To add the linker between the septin Spn1 and fluorescent proteins, standard cloning and gene targeting techniques were used (Bähler *et al*., 1998). Briefly, the linker DNA sequence, flanked upstream by a BamHI site, was added to the 5’ end of the DNA sequence of the fluorophore via a long primer containing the linker DNA sequences and the 5’ sequences of the fluorophore. The DNA sequences of the fluorophores were amplified from pFA6a plasmids (JQW85 for mEGFP, JQW86 for mYFP, and JQW133 for tdTomato) that contained an AscI site directly downstream of the stop codon, and then, they were cloned into the Topo vector. The inserts were released by cutting with BamHI and AscI restriction enzymes and cloned into a pFA6a vector to obtain the templates for gene targeting (JQW221 pFA6a-linker-mEGFP-kanMX6; JQW222 pFA6a-linker-mYFP-kanMX6; and JQW1239 pFA6a-linker-tdTomato-kanMX6). The plasmids were confirmed by sequencing before gene targeting.

### Microscopy and data analyses

Confocal and epifluorescence microscopy were performed as before at ∼23°C with some modifications to minimize perturbation of septin localizations (Davidson *et al*., 2016; Singh *et al*., 2024; Ye *et al*., 2025). Briefly, fresh growing cells from the strains that were woken up on a YE5S plate from -80℃ stocks were inoculated into a 50 ml triple baffled glass flask with ∼10 ml of YE5S liquid medium. The cells were grown in the dark, in a water bath shaker shaking at 150 rpm at 25°C (or 32°C for strains containing the cold-sensitive *cdc4-s16*) for 36 to 48 h. Cells were diluted twice a day with new YE5S medium to ensure that the strains were grown in log phase with OD_595_ < 0.5. Before microscopy, 0.9 ml of culture was mixed with 0.1 ml of 50 μM n-propyl gallate made in YE5S medium and incubated for 2 min to protect cells from free radicals during microscopy. Cells were then collected by gentle centrifugation at 3,000 rpm (845 g) for 30 s. All but ∼10-20 μl of the supernatant was discarded and the cell pellet was resuspended in the remaining medium. ∼3 to 5 μl of the resuspended cells were then spotted onto a gelatin pad (20% gelatin in YE5S liquid medium containing n-propyl gallate at a final concentration of 5 μM) on a microscope slide. The cells were covered with a No. 1.5 cover slip (22 x 22 mm) and sealed with VALAP just before imaging. Cells were imaged with a Nikon CSU-W1 SoRa spinning disk confocal microscope with Hamamatsu ORCA Quest qCMOS camera C15550 on Nikon Eclipse Ti2 microscope with Plan Apo λD 100x/1.45 numerical aperture (NA) oil objective. Yeast mutants or strains with various fluorophores were imaged. For strains with an mYFP/YFP tag, the 514 nm laser was set at 5-20% power; for GFP-based tags, the 488 nm laser at 10-50%; for tdTomato, mScarlet, or other red fluorescent proteins, the 561 nm laser at 10-50% power. For all strains, DIC images associated with fluorescence images were taken to examine cell morphology, but the DIC was somehow compromised by the Piezo used for the confocal imaging.

To observe cells under DIC alone to quantify septation index and cell lysis, cells on the YE5S gelatin pad were observed using a 100×/1.4 numerical aperture (NA) Plan-Apo objective lens on a Nikon Eclipse Ti inverted microscope with a Nikon cooled digital camera DS-QI1 (Nikon, Melville, NY).

Images and fluorescence intensity were analyzed or measured using NIS Elements and Fiji software as previously (Davidson *et al*., 2016; Longo *et al*., 2022; Singh *et al*., 2024). Data in figures and tables are mean ± 1 SD. The *p* values were calculated using two-tailed Student’s t tests.

### Predicting structures of epitope-tagged septins using AlphaFold3

Deep learning artificial intelligence, such as AlphaFold3, has been significantly improved to meet the demands of predicting protein structures and interactions as a complementary tool for experimental data. AlphaFold utilizes its complex neural networks and training strategies to predict the 3D structure of proteins based on their amino acid sequences (Jumper *et al*., 2021; Abramson *et al*., 2024). We developed structural models of septins (Spn1, Spn2, Spn3, and Spn4) through AlphaFold3 (https://alphafoldserver.com/), and predicted their interactions with each other within an octameric structure with different epitope tags. Sequences of each septin protein were obtained through Pombase (https://www.pombase.org/). Each model was run in triplicate and ranked on their predicted template modeling (pTM) score 0-1.0 and interface predicted template modeling (ipTM) score 0-1.0, which are both derived from a template modeling (TM) score that measures the accuracy of the entire structure (Zhang and Skolnick, 2004; Xu and Zhang, 2010). pTM scores that are higher than 0.5 are likely similar to a true structural fold. ipTM scores that are higher than 0.8 are confident predictions, while those under 0.6 are likely to be failures.

Protein sequences were input as quartets in numerical order (for example, Spn1:Spn2:Spn3:Spn4) and doubled for a total of 8 proteins to achieve a predicted octameric structure. Septins Spn1 and Spn4 were attached with various epitope tags at the C-terminus in our predictions. The epitope tag sequences were derived from: 3HA (Addgene plasmid # 39295; http://n2t.net/addgene:39295), tdTomato (Addgene plasmid # 105157; http://n2t.net/addgene:105157), GFP (S65T) (Addgene plasmid # 39292; http://n2t.net/addgene:39292), mEGFP (Addgene plasmid # 105146; http://n2t.net/addgene:105146), and mYFP (Addgene plasmid # 105155; http://n2t.net/addgene:105155) (Bähler *et al*., 1998; Longtine *et al*., 1998; Wu *et al*., 2003; Zhu *et al*., 2013). Each epitope tagged septin contains a small linker sequence (RIPGLIK) between the septin and tag as in the tagged yeast strains (Bähler *et al*., 1998). The highest confidence structures were analyzed using PyMol. Each septin subunit is differentiated by color, and each epitope tag used is further colored distinctly from all septin subunits using PyMol.

## ACKNOWLEDGMENTS

We thank John Pringle, Tom Pollard, Noah DiFilippo, and the former members of the Wu laboratory for fission yeast strains; Nick Ricottilli and Davinder Singh in the lab for technical support; Anthony Vetter from Nikon for microscopy technical support; and current and former members of the Wu lab for helpful discussions and suggestions. We are indebted to John Pringle for sharing unpublished septin immunofluorescence data. J.R.G. and I.M.L. were supported by the Cellular, Molecular, and Biochemical Sciences Training Program (T32 GM141955) and H.E.G. was supported by the Molecular Biophysics Training Program (T32 GM144293) from the National Institutes of Health. The work was supported by the National Institute of General Medical Sciences of the National Institutes of Health (grant R01 GM118746 to J.-Q.W.). The content is solely the responsibility of the authors and does not necessarily represent the official views of the National Institutes of Health.

## Abbreviations

DIC: differential interference contrast
SIN: septation initiation network
SPB: spindle pole body
tdTomato: tandem dimer Tomato
WT: wild type

## SUPPLEMENTAL MATERIALS

## Molecular Biology of the Cell

## Movie Legend

**Movie 1: Comparison of the localization of Spn1-tdTomato and Spn1-linker-tdTomato.** Spn1-tdTomato (JW1345) and Spn1-linker-tdTomato (JW10434) cells were grown in log phase at 25°C for ∼36 h before imaging. The movie shows the maximal intensity projection of 11 focal planes with 0.7 μm spacing at each time point. The images were taken every 1 min for 30 min. 7 frames per second.

---

**Supplemental Table S1:**
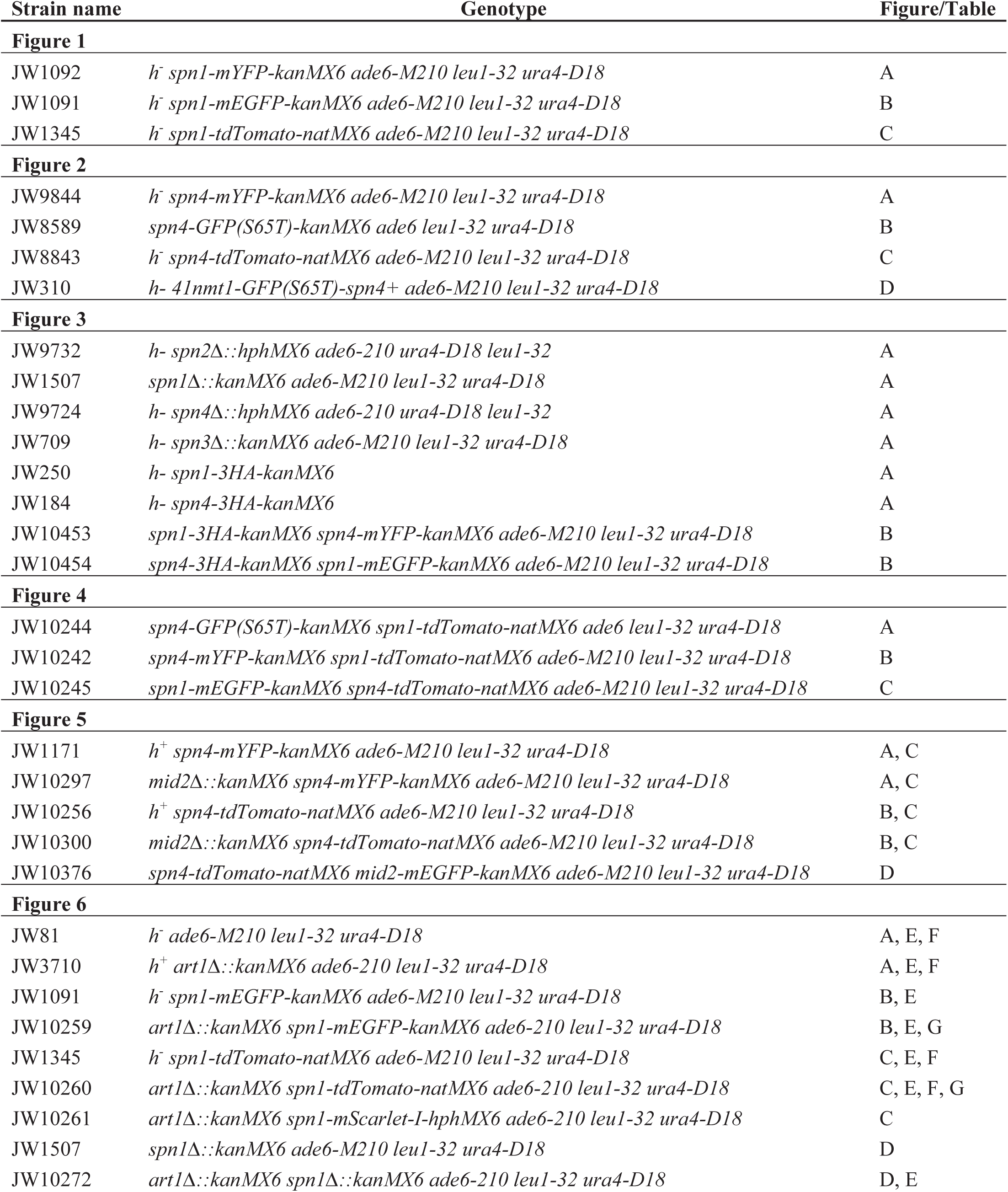
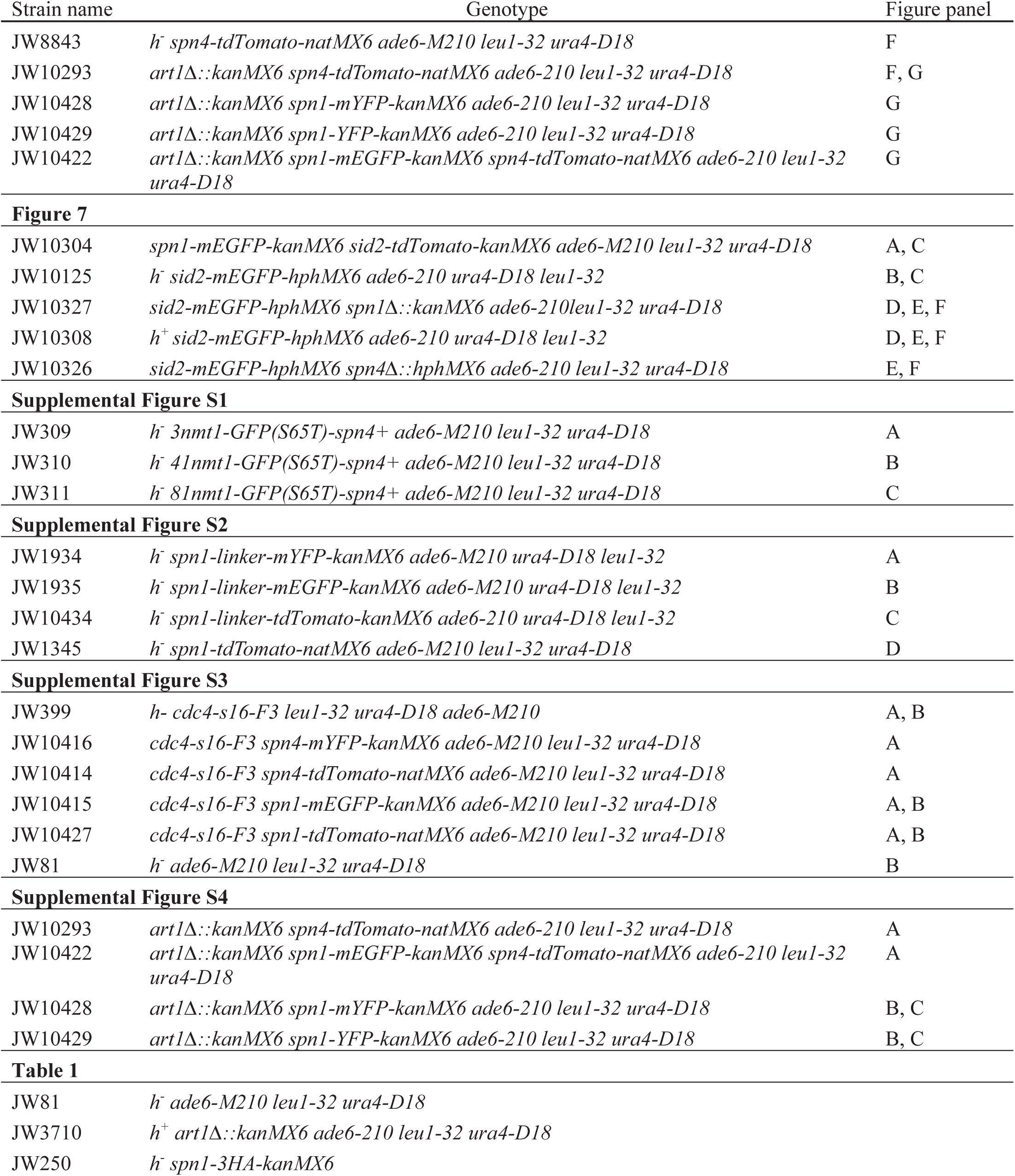
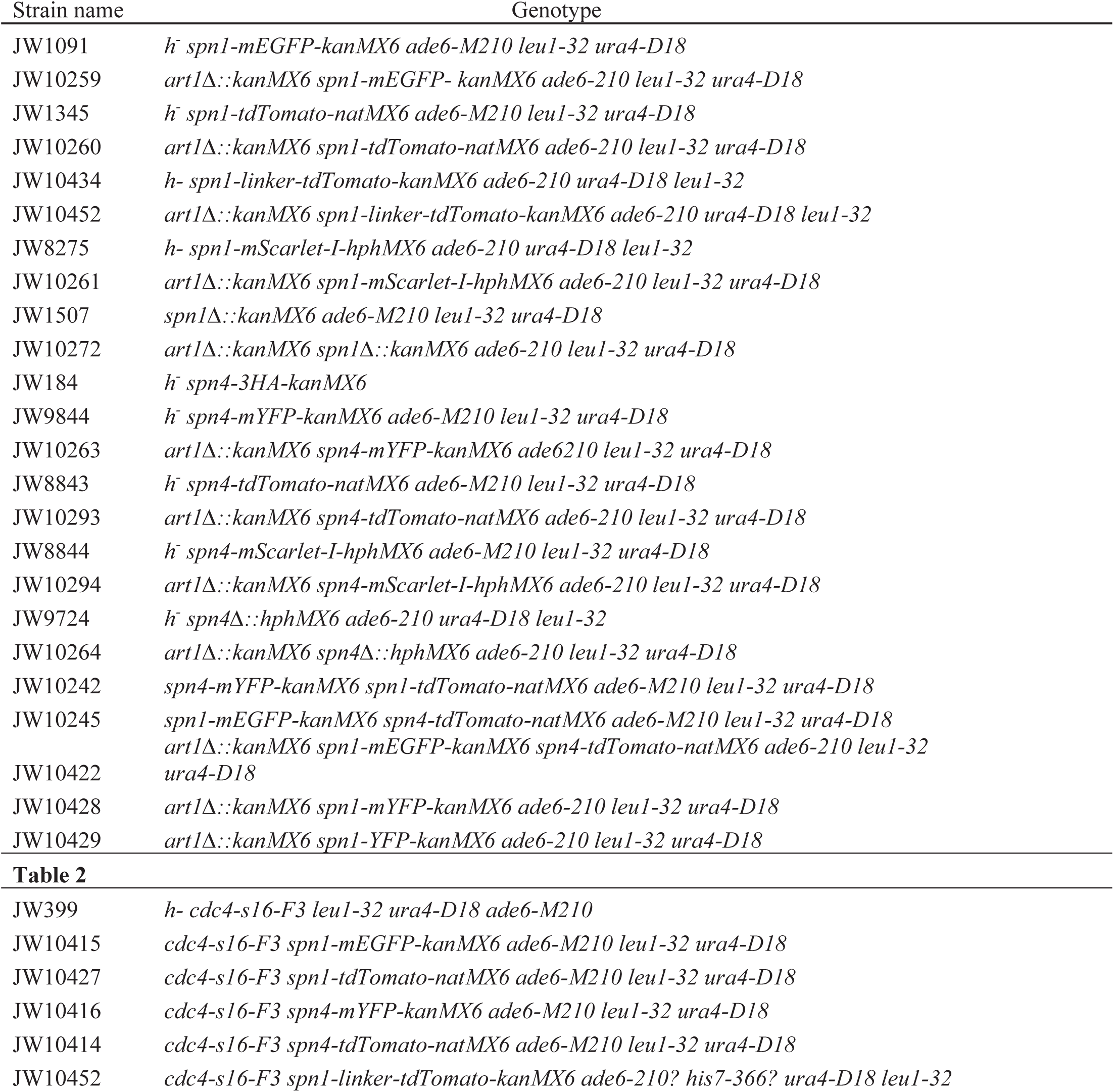
*S. pombe* strains utilized in this study.

**Supplemental Figure S1.**
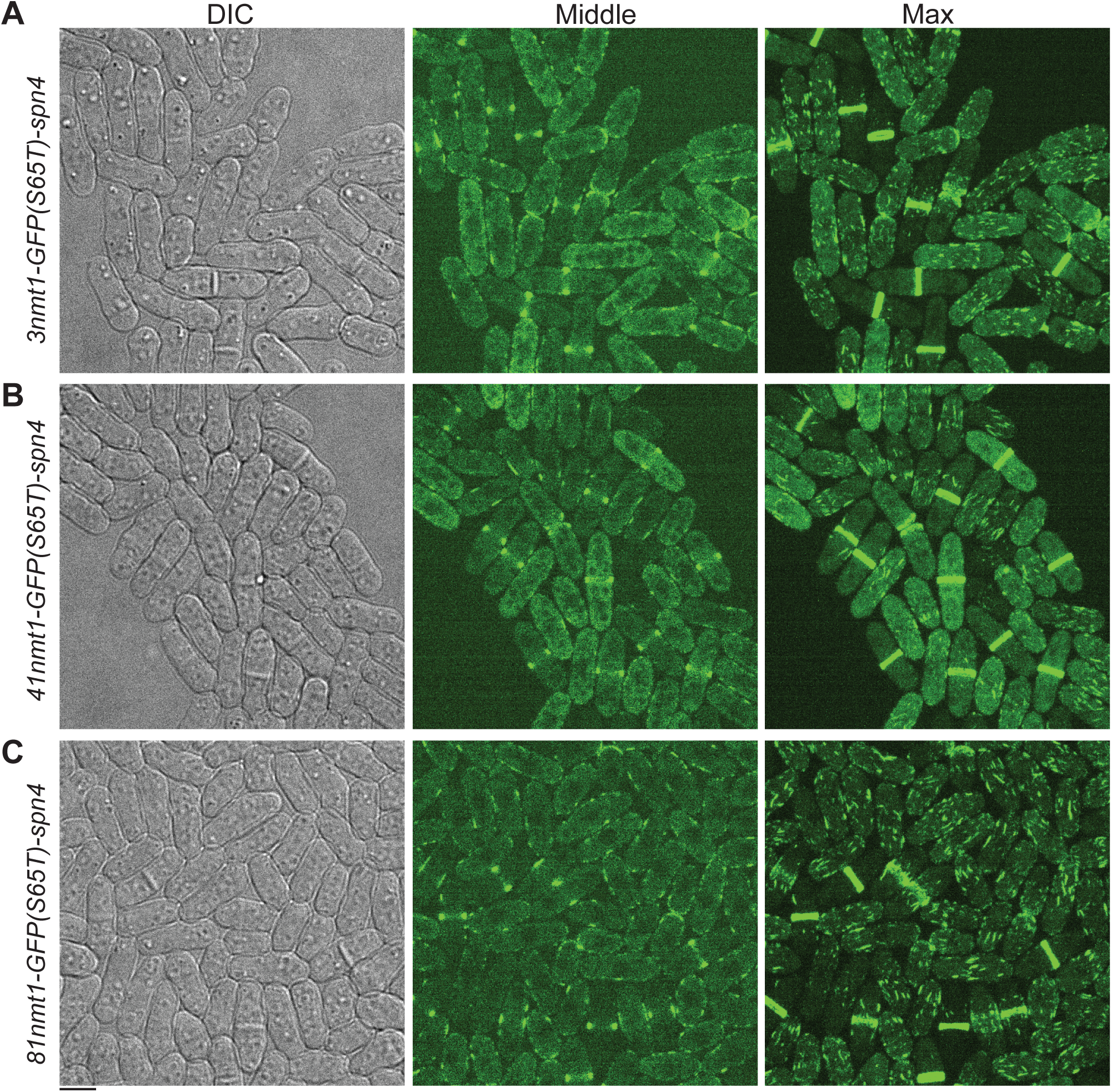
Cells expressing GFP(S65T)-Spn4 at various levels (controlled by the *nmt1* promoters with different strengths) show similar phenotype and localizations. *3nmt1-GFP(S65T)-spn4* cells in YE5S (A), *41nmt1-GFP(S65T)-spn4* cells in EMM5S (B), and *81nmt1-GFP(S65T)-spn4* cells in EMM5S (C). Cells were grown in the indicated liquid media for 20-22 h before imaging. DIC, fluorescence images at the middle focal plane, and the maximal intensity projection of 25 slices with 0.3 µm spacing are shown. Scale bar, 5 µm.

**Supplemental Figure S2.**
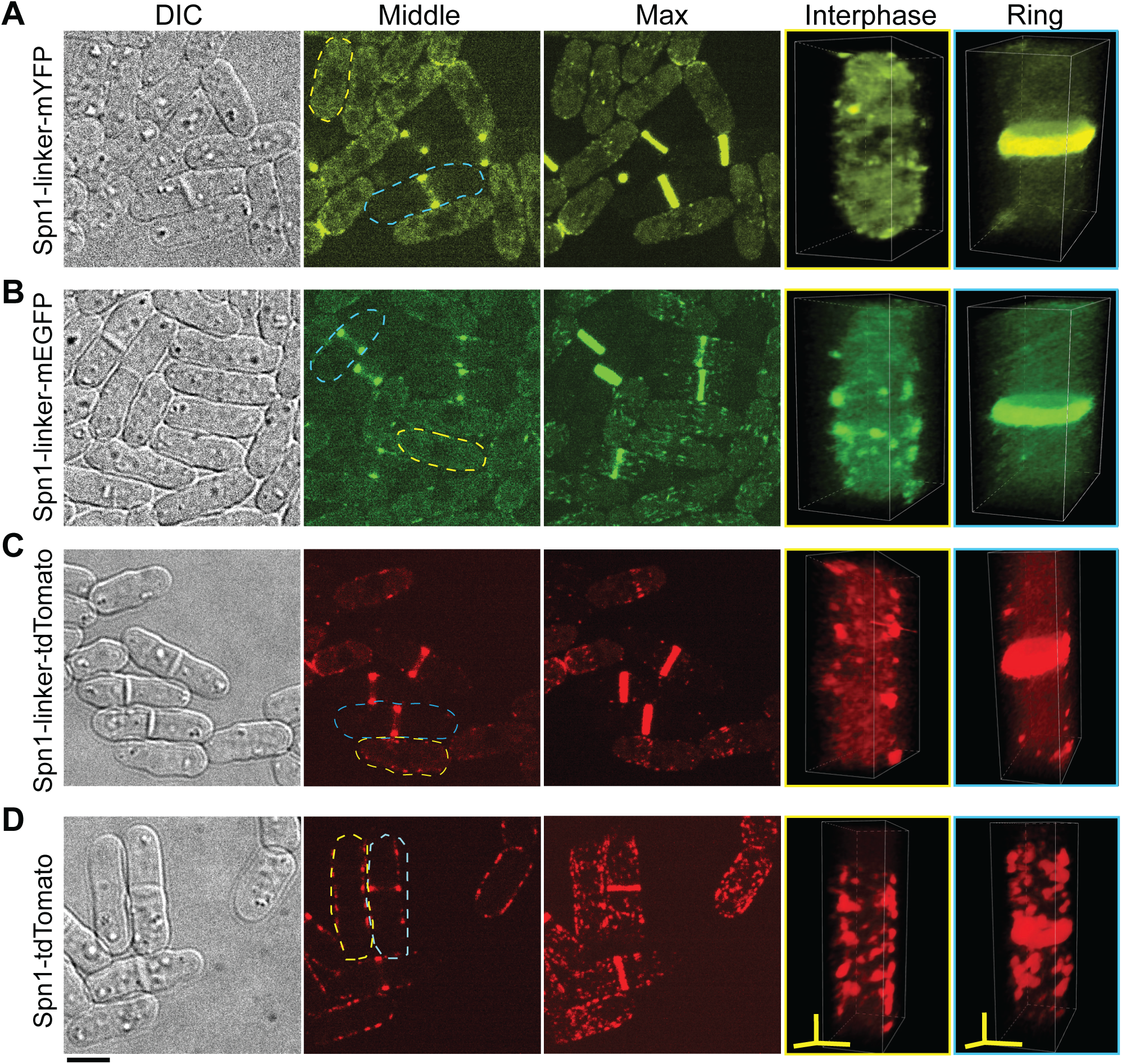
A flexible linker has varying effects on septin Spn1 localizations. Cell morphology and Spn1 localization of cells expressing Spn1-linker-mYFP (A), Spn1-linker-mEGFP (B), Spn1-linker-tdTomato (C), and Spn1-tdTomato cells (D). DIC, fluorescence images at the middle focal plane and the maximal intensity projection are shown. 37 slices with 0.2 µm spacing (A and B) and 19 slices with 0.4 µm spacing (C and D). 3D volumetric projection of representative interphase and dividing cells (Ring) are on the right. Scale bar, 5 µm. Axial bars, 2 µm.

**Supplemental Figure S3.**
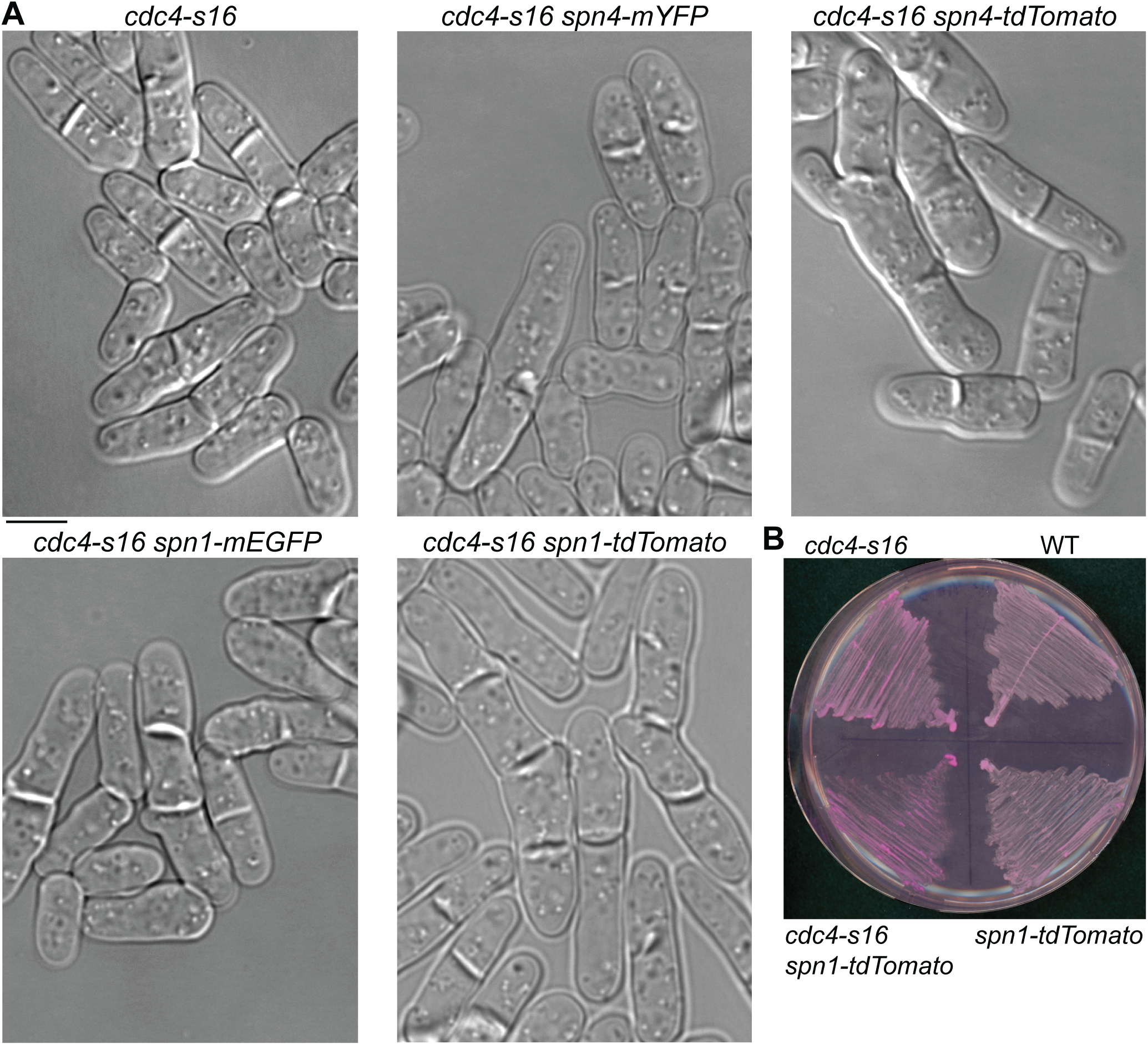
Synthetic genetic interactions between cold-sensitive *cdc4-s16* mutant and tagged septins. (A) Representative DIC images of the indicated strains. Cells were grown in log phase in YE5S liquid medium for ∼36 h at 32°C then shifted to 25°C for 4 h before imaging. Scale bar, 5 µm. (B) Growth of the indicated strains on YE5S + Phloxin B plate at 32°C for ∼29 h.

**Supplemental Figure S4.**
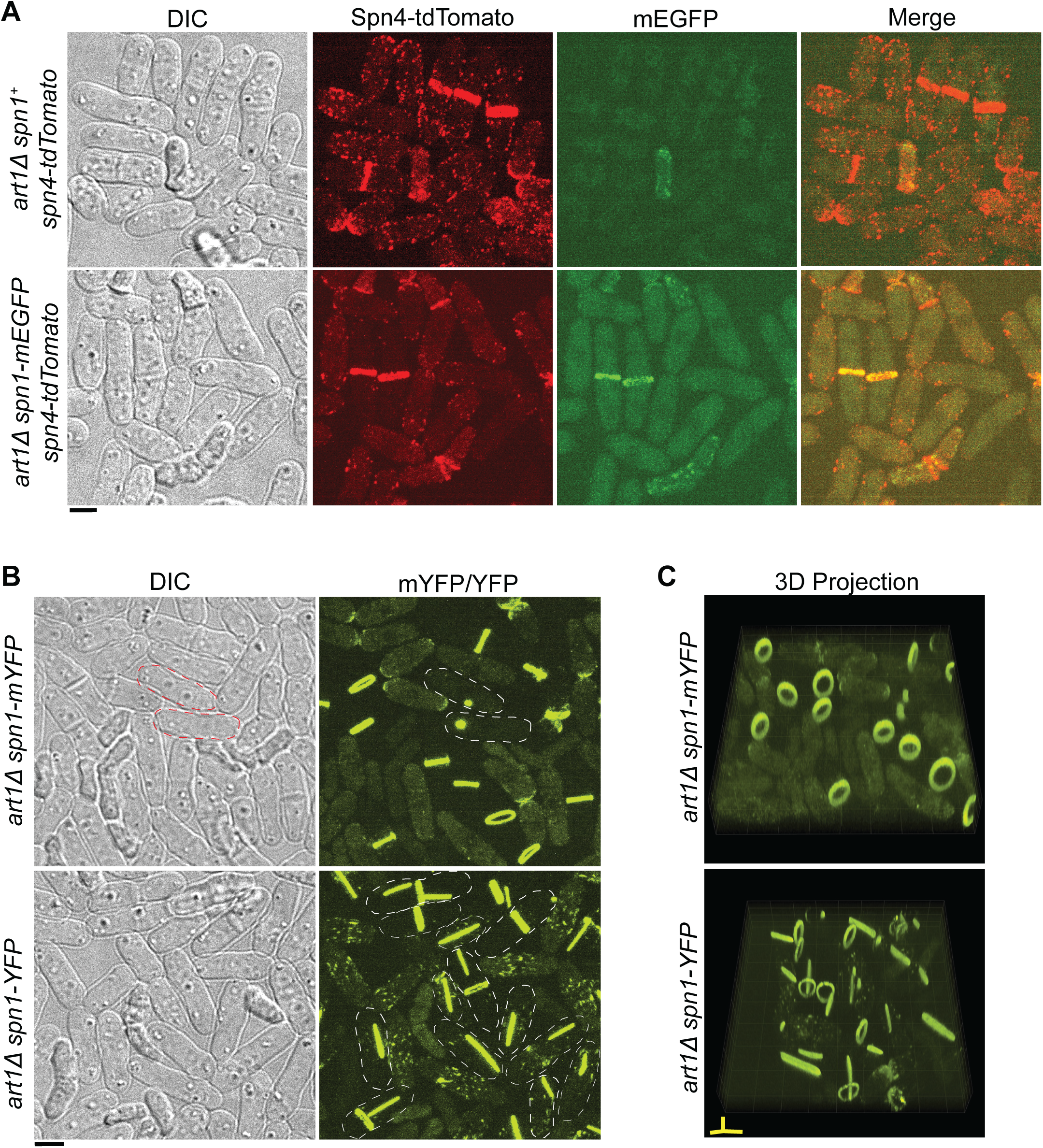
Localizations of the septins Spn1 and Spn4 in *art111* cells. (A) Localizations of Spn4-tdTomato or Spn4-tdTomato Spn1-mEGFP in *art111* cells. Cell morphology and maximal intensity projections of 25 slices with 0.3 µm spacing are shown. Top, untagged Spn1; bottom, Spn1-mEGFP. (B and C) Spn1-YFP but not Spn1-mYFP forms septin bars or spirals in the cytoplasm besides the septin rings at the division site in *art111* cells. (B) DIC and maximal intensity projections of 19 slices with 0.4 µm spacing are shown. (C) 3D projections of fluorescence images. Representative cells are outlined using dashed lines. Spn1-mYFP appears as a discrete spot in a small fraction of cells (outlined). Scale bar, 5 µm.

**Supplemental Figure S5.**
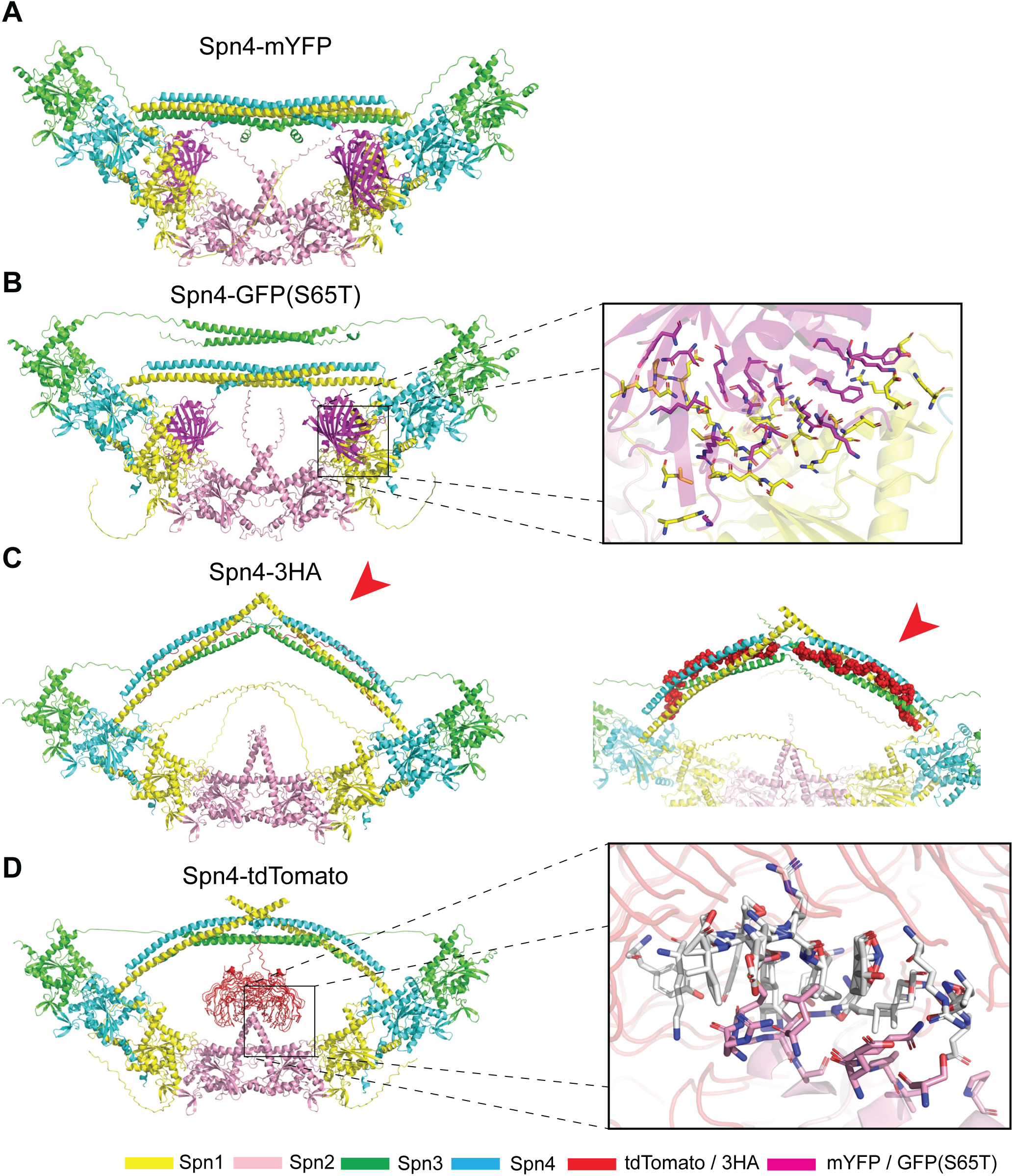
Predicted structures of the septin octameric filament with epitope-tagged Spn4 by AlphaFold3. (A-D) Septins Spn1-4 are differentiated by the color as labeled. Two subunits of each septin were used to construct the octameric filament. tdTomato and 3HA are colored in BR9 (red), and GFP(S65T) and mYFP are colored in magenta. Red arrowheads indicate the 3HA interaction with the bundled coiled coils. (B and D) Box projections show interacting residues between (B) GFP(S65T) (magenta) and Spn1 (yellow) or (D) tdTomato (white) and Spn2 (pink) within 3.5Å, indicative of Van der Waals forces represented as sticks. (A) Spn4-mYFP model, pTM = 0.45. (B) Spn4-GFP(S65T) model, pTM = 0.45. (C) Spn4-3HA, pTM = 0.49. (D) Spn4-tdTomato model, pTM = 0.47.

**Supplemental Figure S6.**
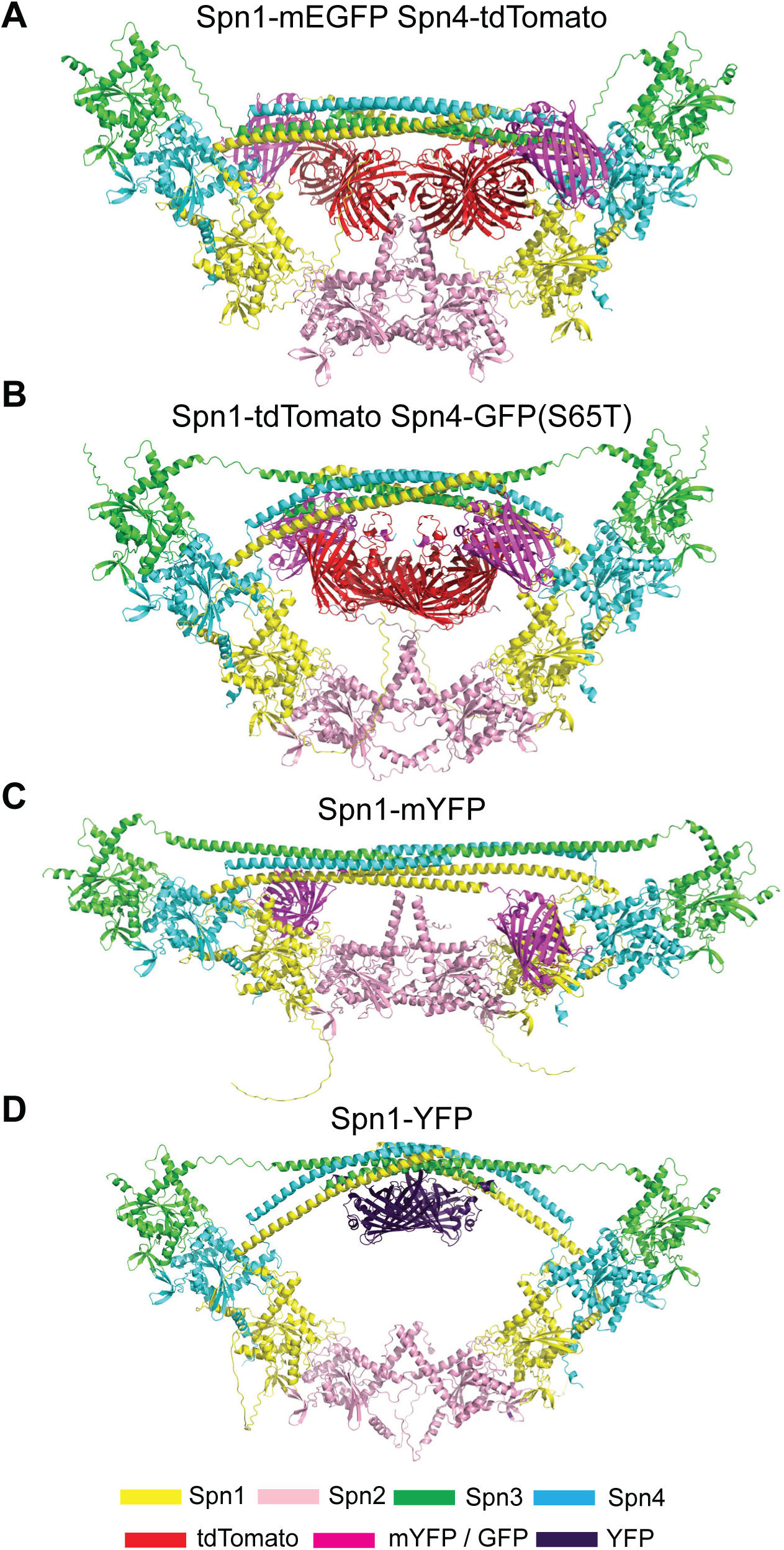
Predicted structures of the septin octameric filament of monomeric and dimeric YFP-tagged Spn1 and the cooperative colocalization effect by AlphaFold3. (A-D) Septins Spn1-4 are differentiated by the color as labeled. Two subunits of each septin were used to construct the octameric filament. tdTomato is colored in BR9 (red), GFP/mYFP is colored in magenta, and the dimeric YFP is colored in francium (dark purple). (A) Spn1-mEGFP Spn4-tdTomato model, pTM = 0.41. (B) Spn1-tdTomato Spn4-GFP(S65T) model, pTM = 0.42. (C) Spn1-mYFP model, pTM = 0.49. (D) Spn1-YFP model. pTM = 0.45.

